# Bioactive coatings on 3D printed scaffolds for bone regeneration: Use of Laponite^®^ to deliver BMP-2 in an ovine femoral condyle defect model

**DOI:** 10.1101/2024.02.25.581921

**Authors:** Karen M. Marshall, Jane S. McLaren, Jonathan P. Wojciechowski, Sebastien J. P. Callens, Cécile Echalier, Janos M. Kanczler, Felicity R. A. J. Rose, Molly M. Stevens, Jonathan I. Dawson, Richard O.C. Oreffo

**Author notes:** Corresponding Authors: Dr Karen Marshall and Professor Richard OC Oreffo. Joint first authors (Karen M. Marshall, Jane S. McLaren).

## Abstract

Biomaterial-based approaches for bone regeneration seek to explore alternative strategies to repair non-healing fractures and critical-sized bone defects. Fracture non-union occurs due to a number of factors resulting in the formation of bone defects. Rigorous evaluation of the biomaterials in relevant models and assessment of their potential to translate towards clinical use is vital. Large animal experimentation can be used to model fracture non-union while scaling-up materials for clinical use. Growth factors modulate cell phenotype, behaviour and initiate signalling pathways leading to changes in matrix deposition and tissue formation. Bone morphogenetic protein-2 (BMP-2) is a potent osteogenic growth factor, with a rapid clearance time *in vivo* necessitating clinical use at a high dose, with potential deleterious side-effects. The current studies have examined the potential for Laponite^®^ nanoclay coated poly(caprolactone) trimethacrylate (PCL-TMA900) scaffolds to bind BMP-2 for enhanced osteoinduction in a large animal critical-sized bone defect. An ovine femoral condyle defect model confirmed PCL-TMA900 scaffolds coated with Laponite^®^/BMP-2 produced significant bone formation compared to the uncoated PCL-TMA 900 scaffold *in vivo*, assessed by micro-computed tomography (µCT) and histology. This indicated the ability of Laponite^®^ to deliver the bioactive BMP-2 on the PCL-TMA900 scaffold. Bone formed around the Laponite^®^/BMP-2 coated PCL-TMA900 scaffold, with no erroneous bone formation observed away from the scaffold material confirming localisation of BMP-2 delivery. The current studies demonstrate the ability of a nanoclay to localise and deliver bioactive BMP-2 within a tailored octet-truss scaffold for efficacious bone defect repair in a large animal model with significant implications for translation to the clinic.

## 1. Introduction

Advances in healthcare have resulted in a welcome increase in global longevity although this is accompanied by additional healthcare challenges, including the prevalence of chronic disease and age-related tissue degeneration. Tissues can fail to heal after injury, such as bone following a fracture. Bone defects can arise from a variety of causes and amputation may be required, due to extensive soft tissue, vascular and/or nerve damage, making the limb unsalvageable (1, 2). Advancements in tissue engineering have enabled the development of materials that resemble, in part, the composition and functionality of the damaged tissue, thereby reducing the need for grafting or transplantation procedures. Grafting of tissue has inherent risks, including the potential for immune-mediated rejection or infection. Bone tissue engineering seeks to develop materials to substitute the patient’s bone, autograft, to encourage healing of the bone defect (3, 4). Hence, the ideal bone substitute material is biocompatible, biodegradable, osteoconductive, osteoinductive, porous, strong, easy to use and cost effective (1).

BMP-2 is a clinically proven and acknowledged potent osteoinductive factor although, BMP-2 is commonly administered clinically at a high concentration in humans to elicit a biological response due to the short half-life of BMP-2 *in vivo*. The BMP-2 is delivered in a collagen sponge carrier by adding a set volume of BMP-2 solution to be absorbed. These high concentrations of BMP-2 may saturate the binding sites of the scaffold and allow desorption and release of growth factor in a rapid burst upon implantation (5). Therefore, BMP-2 has been immobilised onto scaffolds, to maintain bioactivity while minimising potential side-effects, using various methods including BMP-2 binding to avidin-biotin, fibronectin, or heparin or within nanoparticles, hydrogels, or microspheres (6–12).

Laponite^®^ (herein Laponite) is a biocompatible and biodegradable synthetic smectite clay with the formula (Na_0.7_Si_8_Mg_5.5_Li_0.3_O_20_(OH)_4_). When dispersed in water, Laponite forms a colloidal dispersion of charged disk-shaped nanoparticles, of thickness ca 1 nm and diameter ca 25 nm which at concentrations above 2% w/w, self-assemble to form a reversible gel (13). Clays have historically found use in pharmaceuticals and cosmetics, as sorbents and rheology modifiers, and have recently received interest in tissue engineering and regenerative medicine due to their ability bind and localise proteins. Laponite has previously been shown to sequester BMP-2 to induce osteogenic differentiation of skeletal cells and to be efficacious in the delivery BMP-2 in murine subcutaneous implantation and murine femur defect studies (14–17).

A limitation of small animal models is that the healing rates are faster in rodents and rabbits than in humans, therefore large animal models such as sheep, goats and dogs may provide models more comparable to the clinical scenario (18). Larger animals have been used in bone tissue engineering research to allow implantation of larger scaffolds and increased biomechanical challenge to be assessed, mimicking the clinical situation. To that end, sheep have been widely used as an experimental model for bone tissue engineering, including osteoporosis research (19–22).

The current work details a 3D-printed robust octet-truss design scaffold composed of an optimised formulation of poly(caprolactone) trimethacrylate (PCL-TMA900). This polymer permitted the printing of intricate shapes with a reduced risk of brittle failure, as the polymer was more compliant and less brittle than the lower molecular weight PCL trimethacrylate (PCL-TMA) used in our previous study (23).

The overall aim of this study was to investigate the use of Laponite with BMP-2 as a dry, ready-to-use, biocompatible, bioactive, reliable, osteoinductive coating on PCL-TMA900 scaffolds for bone tissue engineering in an ovine critical-sized bone defect model. This required retention of active BMP-2 on/in the Laponite surrounding the PCL-TMA900 scaffold, in sufficient quantities to induce osteogenic differentiation of skeletal cell populations to produce significant bone volumes *in vivo*, augmenting bone repair compared to an uncoated PCL-TMA900 scaffold (16, 24, 25). Given the biocompatibility of Laponite and BMP-2, the Laponite/BMP-2 coating could, it was proposed, easily and practically be applied to induce bone formation around a large PCL-TMA900 scaffold *in vivo* (14–17, 20). The PCL-TMA900 was printed in an octet-truss scaffold design, with Laponite adherence and sites of preference on the scaffold determined by addition of barium prior to micro-computed tomography (µCT) imaging of the scaffolds. The retention and delivery of BMP-2 at physiologically active concentrations was investigated to ensure the potential for bone formation by the scaffold prior to implantation. Finally, following *in vitro* confirmation of the ability of the coating to adhere to the scaffold and, from previous *in vivo* studies illustrating induced bone formation, the PCL-TMA900 scaffolds coated with Laponite and BMP-2 were applied to an ovine femoral condyle defect model (14, 17). This large animal model of bone tissue engineering was performed with a view to inform the potential for clinical translation of the Laponite and BMP-2 coated PCL-TMA 900 material. The *in vivo* ovine critical-sized femoral defect study employed a bilateral, cylindrical bone defect of 8 mm diameter and 15 mm depth in the metaphysis of the medial femoral condyles, with 8-10 mm diameter and 13-20 mm depth previously reported in sheep (26). The femoral condyles were harvested after 13 weeks for assessment by µCT and histology to determine the efficacy of the Laponite/BMP-2 coating on the PCL-TMA900 scaffolds to induce bone formation critical in any subsequent safety testing and regulatory approval on the path towards clinical translation.

## 2. Materials and Methods

### 2.1 Materials

Reagents were purchased as follows: Recombinant human BMP-2 (Infuse/InductOS Bone graft kit, Medtronic, USA); BMP-2 Quantikine ELISA Kit (biotechne^®^, R&D systems, UK); Alcian blue 8X, Light green SF, Orange G 85% pure, Paraformaldehyde 96% extra pure, phosphomolybdic acid hydrate 80% (Acros Organics); Picrosirius Red, Van Gieson’s stain, Weigert’s Haematoxylin Parts 1 and 2 (Clintech Ltd, UK); PBS, trypsin/EDTA, DMEM, Penicillin-Streptomycin (Scientific Laboratory Supplies, SLS); 4-nitrophenol solution 10 nM, acetic acid, acetone, acid fushsin, alizarin red S, fast violet B salts, histowax, Naphthol AS-MX phosphate 0.25%, ponceau xylidine, silver nitrate, sodium hydroxide pellets (Merck, UK); dibutyl phthalate xylene (DPX), fetal bovine serum, histoclear (Thermofisher Scientific, UK); fast green and sodium thiosulphate (VWR); Isoflurane (Dechra, UK), and Vetasept^®^ were sourced from Cockburn Vets, 100 London Road, Coalville, UK. Size 1 and 0 Vicryl suture (Ethicon, USA) and Fentanyl, Elastoplast and bandages, medetomidine, ketamine, propofol, rimadyl, lidocaine, from Cockburn Vets, 100 London Road, Coalville, UK. All other consumables and reagents were purchased from Sigma-Aldrich, UK. Poly(caprolactone) triol, trimethylolpropane initiated, M_w_ = 830 Da was purchased from BOC Sciences, UK.

### 2.2 Production and printing of PCL trimethacrylate scaffold material

#### 2.2.1 Poly(caprolactone) trimethacrylate synthesis

The synthesis of PCL-trimethacrylate of this molecular weight and 3D-printing using masked stereolithography (mSLA) has been previously reported (27–29). As previously detailed in Marshall *et al*. (14, 17), to synthesise the PCL-TMA900 material, poly(caprolactone) triol, M_n_ = 830 Da, (100 g, 0.12 mmol, 1 eq), anhydrous dichloromethane (300 mL) and triethylamine (100 mL, 0.72 mmol, 6 eq) were added to a 1 L two-necked round bottom flask. The reaction was placed under a nitrogen atmosphere and then cooled in an ice-water bath for 15 minutes. A pressure-equalising dropper funnel charged with methacryloyl chloride (53 mL, 0.67 mmol, 4 eq) was attached to the round bottom flask. The methacryloyl chloride was added dropwise over approximately 3 hours. The reaction was covered with aluminium foil to protect it from light and allowed to stir and warm to room temperature (RT) overnight. The following day, methanol (50 mL) was added to quench the reaction, which was allowed to stir at RT for 30 minutes. The reaction mixture was dissolved in dichloromethane (800 mL), transferred to a separating funnel and washed with 1 M aqueous hydrochloric acid solution (5 × 250 mL), saturated sodium bicarbonate solution (4 × 400 mL) and brine (1 × 400 mL). The organic layer was then dried with anhydrous magnesium sulphate, filtered and concentrated via rotary evaporation. The crude yellow liquid was then purified using a silica plug, with dichloromethane as the eluent. Fractions containing PCL-trimethacrylate were pooled and concentrated via rotary evaporation. The PCL-trimethacrylate was transferred to a brown glass vial and dried using a stream of air (through a plug of CaCl_2_) overnight to yield the title compound as a slightly yellow viscous liquid (94.25 g). The PCL-trimethacrylate was supplemented with 200 ppm (w/w) of 4-methoxyphenol (MEHQ) as an inhibitor.

^1^H NMR (400 MHz, CDCl_3_) δ 6.11 – 6.05 (m, 3H), 5.61 – 5.50 (m, 3H), 4.16 – 3.99 (m, 18H), 2.34-2.27 (m, 12H), 1.96 – 1.88 (m, 9H), 1.71 – 1.60 (m, 26H), 1.47 – 1.31 (m, 12H), 0.98 – 0.83 (m, 3H).

The characterisation data agrees well with that previously reported (27). The degree of functionalisation was determined as >90% (14).

#### 2.3.2 Scaffold design and 3D printing of PCL-trimethylacrylate scaffolds

The octet-truss scaffold shape was designed using a custom Matlab script to fit a cylindrical domain with 8 mm diameter and 15 mm height. The cubic unit cell for the octet-truss lattice was designed to have a width, depth, and height of 3.8 mm and a strut diameter of approximately 1.1 mm. The scaffolds had a surface area of 1382.3 mm^2^ and a relative density of 56%. The PCL-TMA900 scaffolds were printed using mSLA 3D printing on a Prusa SL1S (**Supplementary Figure 1**). The resin was prepared for 3D printing by first dissolving 0.1% (w/w) 2,5-thiophenediylbis(5-tert-butyl-1,3-benzoxazole) (OB+) as a photoabsorber in the PCL trimethacrylate by stirring at RT for 1 hour. Finally, 1.0% (w/w) diphenyl(2,4,6-trimethylbenzoyl)phosphine oxide (TPO-L) as a photoinitiator was added to the resin. The scaffolds were sliced in PrusaSlicer with their longitudinal axis parallel to the surface of the build platform. Supports and pads were used to raise the objects off the build platform, with ten supports placed along the scaffold to ensure adhesion to the build platform during the printing whilst not compromising the structure. The scaffolds were sliced at a layer height of 75 µm using a 25 second exposure time per layer.

After printing, the scaffolds were rinsed in ethanol and removed from the build plate. Scaffolds were sonicated in ethanol (5 × 5 minutes) and allowed to dry for 15 minutes at RT. The scaffolds were post-cured using a Formlabs Form Cure for 60 minutes at RT. After post-curing, the scaffolds were soaked into ethanol overnight at RT on a rocker (100 rpm), rinsed with ethanol (3 ×) and allowed to dry at RT before being EO sterilised.

### 2.3 Laponite (1% w/w) production

1% (w/w) Laponite was made 48 hours prior to use by the addition of 0.1 g Laponite (BYK-Altana) to 9.9 g dH_2_O slowly in a glass jar, while mixing at 250 rpm with a magnetic stirrer at RT and left to stir for 1-2 hours to ensure complete dissolution. The jar was weighed and autoclaved at 126°C in a bench-top autoclave and reweighed. In a class II microbiological safety cabinet (MSC), the mass of dH_2_O lost was replaced by adding sterilised dH_2_O until the original mass was reached and stored at 4°C.

### 2.4 Laponite and BMP-2 coating of scaffolds

The PCL-TMA900 scaffolds (8 mm wide x 15 mm height) were placed individually in 2 mL 1% Laponite in a 15 mL falcon tube for 1 hour at RT. The scaffolds were then removed and left to dry individually on 25-gauge needles held by sterile bone wax (Ethicon, UK) to the lid of 50 mL falcon tubes for 5 hours. Once dry, the scaffolds were immersed in 1.5 mL BMP-2 (100 µg/mL) diluted in PBS in a low protein-binding falcon tube (Eppendorf, UK). The bottom of the tube was gently tapped on a surface to remove any bubbles between the pores of the scaffold to allow the solution to completely contact the scaffold. The scaffolds were left at RT for 24 hours prior to removal from the Eppendorf tube and were left to dry overnight on 25-gauge needles, prior to the 50 mL tube being screwed on to the lid, keeping the needle pointing upwards to keep the scaffolds in a sterile environment (**Supplementary Figure 2**). Prior to the *in vivo* study, Laponite was confirmed to adhere and persist on the scaffold material after immersion and agitation in dH_2_O (**Supplementary information Figure 3**).

An ELISA was performed with a Quantikine^®^ ELISA BMP-2 immunoassay kit using the BMP-2 solutions remaining from optimisation experiments involving the BMP-2 (100 µg/mL or 200 µg/mL) coating process; to determine the quantity of BMP-2 adhered to each Laponite coated PCL-TMA900 scaffold (**Supplementary information Figure 4 and Figure 5**). Prior to the *in vivo* study, the optimal concentration of BMP-2 was confirmed on the scaffold material when 100 µg/mL was used and the BMP-2 was confirmed to also be released at a bioactive concentration by use of ALP staining of C2C12 cells exposed to media in which the scaffolds had been incubated (**Supplementary information Figure 6**). Further, an ELISA was performed on the BMP-2 solutions (100 µg/mL initial concentration) remaining from coating the scaffolds implanted into the sheep to determine the average mass of BMP-2 on the scaffolds (**Supplementary information Figure 7**).

### 2.5 *In vivo* sheep femoral defect study

#### 2.5.1 Sheep information and housing

All procedures were performed in accordance with institutional guidelines, with ethical approval and under project license (PPL) PP9657326, in accordance with the regulations in the Animals (Scientific Procedures) Act 1986. Further, the study used the ARRIVE guidelines and was approved by the University of Nottingham Animal Welfare and Ethical Review Body (AWERB) and University of Southampton Ethics and Research Governance Online (ERGO II) committee. Sixteen, skeletally mature, non-pregnant, adult (>2 years) English mule ewes (average weight 74.3 ± 6.1 kg, average body condition score (BCS) 2.90 ± 0.17) were randomly assigned to four treatment groups (**Table 1**). It should be noted that each sheep received a different treatment in each limb (**Supplementary Figure 8)**. Sheep were group housed in a barn and had access to *ad libitum* hay and water for the duration of the study. Animals were examined to ensure good physical condition and were acclimatised to the indoor barn environment for a minimum of 7 days prior to surgery.

**Table 1:** Groups of sheep allocated in the study. One sheep with autograft and uncoated PCL-TMA900 scaffold implants died <24 hours post-operatively due to unknown causes as per the post-mortem report, *unrelated* to the surgical procedure or scaffold implantation.

| Group | Purpose | Number of defects |
| --- | --- | --- |
| Empty defect | Control group | 8 |
| Autograft | Control group | 7 |
| Uncoated PCL-TMA900 scaffold | Comparator group | 7 |
| Laponite/BMP-2 coated PCL-TMA900 scaffold | Comparator group | 8 |

#### 2.5.2 Implant preparation

There were 4 groups in the study; i) empty defect control to show non-union of the bone defect over the 13 week study period; ii) autograft, comprised of 2 cores of bone harvested at surgery as the positive control, to show reliable healing of the bone defect; iii) uncoated PCL-TMA900 scaffolds as the negative control scaffold group; and, iv) Laponite/BMP-2 coated scaffolds as the test scaffold group with n=8 defects initially per group.

In preparation for the sheep study, to assess the size of the scaffold for fit testing, an 8 mm diameter, 15 mm deep defect was created in the medial femoral condyle and the 3D printed 8 mm diameter, 15 mm deep scaffold inserted, allowing press-fit of the scaffold to ensure the scaffold would be secure and simple to implant (**Supplementary Figure 9)**.

#### 2.5.3 Medial femoral condyle defect surgical procedure

The method is extrapolated from M^c^Laren *et al*. (30). The day prior to surgery, the neck was clipped over a jugular vein and the skin of the lateral antebrachium of one forelimb was clipped, cleaned and de-greased prior to application of Fentanyl cutaneous patches to deliver approximately 2 µg/kg/hr Fentanyl (Durogesic patches, Janssen-Cilag, Saunderton, UK) held by Elastoplast (Beiersdorf, Australia) and a 2^nd^ bandage layer (Easifix^®^ Cohesive, Leukoplast^®^) to provide pre-operative and up to 3 days post-operative analgesia.

Prior to surgery food was withheld for a minimum of 12 hours, and water was withheld for ∼2 hours prior to surgery to reduce the risk of regurgitation and aspiration. Thirty minutes prior to surgery, local anaesthetic cream (Emla 5% cream, Aspen Pharmacare, UK) was applied topically to the clipped region of the neck. The neck was then prepared with chlorhexidine scrub (Hibiscrub, Mölnlycke) and ethycalm (Invicta Animal Health) was used to anaesthetise the skin prior to insertion of a 16 gauge 3.25 inch intravenous catheter (BD Angiocath). Induction of anaesthesia using Ketaset (Ketamine, 5 mg/kg IV, Fort Dodge Animal Health Ltd, Southampton, UK) and 0.3 mg/kg Hypnovel (Midazolam, Roche Products, Welwyn Garden City, UK) was performed. The larynx was sprayed with local anaesthetic (Intubease^®^, Dechra, UK) and after one minute to allow the solution to take effect, a single dose of Alfaxan (Alfaxan multidose, 10 mg/mL, 2 mL IV, Jurox) was administered and intubation performed. Ophthalmic ointment (GelTears 0.25% carbomer eye gel) was applied to both eyes. Anaesthesia was maintained with ∼2 % Isoflurane (Isoflurane-Vet, Merial, UK) in 100 % oxygen. Pre- and 2 days post-operative analgesia was given by means of carprofen (Rimadyl™ 4 mg/kg, 50 mg/mL, Zoetis) intra-muscular (IM) injection. Prophylactic antibiotics (Pen & Strep Suspension for Injection, 0.04 mL/kg, Norbrook) were given pre- and post-operatively the following day. For surgery, the animals were placed in dorsal recumbency to access the medial aspect of both hind legs. The stifle area was then clipped and prepared using chlorhexidine scrub solution (Hibiscrub, Mölnlycke) and 5 mL local anaesthetic bupivacaine hydrochloride (Marcaine, Pfizer, USA) injected around the incision site. Anti-microbial gel (Virusan^®^ Gel, amity) was then applied and left to dry. Once in theatre, the area was prepared with a single use chlorhexidine/isopropyl alcohol applicator (BD ChloraPrep 3 mL Applicator 2% w/v/70% Cutaneous Solution). The skin and underlying soft tissue over the medial femoral condyle were incised and a single cylindrical drill hole/defect (8 mm diameter × 15 mm deep) was created in the cancellous bone region of the medial femoral condyle. The defect was cleared of debris using a custom calibrated reamer to create a standardised defect. Throughout coring and reaming, the drills were cooled with sterile saline solution to prevent tissue damage. The defect was lavaged with sterile saline to remove any bone fragments and dried using a sterile swab to check for bleeding, however none/very minor ooze was noted. For implantation, the uncoated PCL-TMA900 scaffold or Laponite/BMP-2 coated PCL-TMA900 scaffold was inserted into the bone defect using digital pressure or forceps, respectively. Defects left empty or filled with autograft were used as negative and positive controls, respectively. The empty defect had nothing implanted, while autograft was formed from 2 bone cylinders produced from drilling out the defects to produce enough bone to fill one defect. The autograft with removal of the cortical bone/periosteum was macerated into small cancellous bone fragments using rongeurs, prior to implantation into the defect site. After implantation, the periosteum was sutured over each defect/scaffold with polyglactin 910 suture (Vicryl size 1, Ethicon, Kirkton, UK) which aided retention of autograft in the bone defect. The subcutaneous tissue was closed using absorbable material (Vicryl size 1, Ethicon, Kirkton, UK) in simple continuous suture pattern, whilst the skin was closed with absorbable suture material (Vicryl size 0, Ethicon, Kirkton, UK) in an intradermal, simple continuous suture pattern. A moisture vapour-permeable spray dressing (KRUSSE Wound Plast Spray, Denmark) was then applied over the wound for protection from contamination. Images of the surgical procedure are shown in **Supplementary Figure 10**. After surgery, animals were housed overnight in an indoor pen with the other sheep who had surgery on the same day, prior to returning to group housing (**Supplementary Figure 11**). Daily grimace and mobility scoring was performed to monitor for any signs of pain or complications.

#### 2.5.4 Sheep aftercare, tetracycline labelling of bone and study end

The sheep were injected with oxytetracycline (Terramycin LA, 200 mg/ml, Pfizer) at weeks 6 and 11 post-operatively to label new bone deposition to allow visualisation with fluorescence microscopy. The sheep were closely monitored postoperatively for any clinical signs of pain using lameness and grimace scoring. Throughout the study, body weight and body condition scores were recorded for all sheep. Body condition scores highlighted the health and amount of fat on an animal, with a score of 1 indicating an emaciated animal and a score of 5 indicating an obese animal, with a score of 3 classed as optimal. Animals were sacrificed after 13 weeks post-operatively by intravenous barbiturate (Somulose, Dechra) overdose. The femoral condyles were retrieved immediately, and fixation initiated at RT in 10% neutral buffered formalin (NBF, Sigma-Aldrich), followed by fixation/storage at 4°C. The bones were trimmed for Micro-CT (µCT) and histological analysis.

#### 2.5.5 Micro-CT procedure and analysis of results

Micro-CT was performed using a MILabs OI-CTUHXR preclinical imaging scanner (Utrecht, The Netherlands) to determine bone ingrowth within the femoral condyle defects. Micro-CT reconstructions were obtained via MILabs software (MILabs-Recon v. 11.00). Imaging of *ex vivo* bone at week 13 was performed (computer settings detailed in **Supplementary Information Table 1**). The condyle was initially trimmed to remove the lateral condyle to enable the bone to fit in the µCT scanner for image reconstruction at 40 µm (**Supplementary Figure 12 a and b**). Further trimming to obtain a smaller volume of bone to obtain higher resolution images at 20 µm was performed (**Supplementary Figure 12 c**). An approximately 2.5 cm × 2.5 cm wide × 4 cm deep cuboidal volume of bone in the region of interest around the defect was selected using a fine-toothed hacksaw blade and hacksaw (Wickes, UK), with the condyle held in a vice (Stanley, UK) for trimming. A density phantom, as a reference for quantification of bone density, was scanned using the same acquisition and reconstruction parameters at 40 µm and 20 µm. Example µCT images of the defect in a cadaver sheep are shown in **Supplementary Figure 13** to show the defect size and position. Formation of bone was assessed using Imalytics Preclinical software v3.0 (Gremse-IT GmbH). A gauss filter of 1.5 and bone density threshold for analysis was set equal to the average result from the lower density bone phantom. A cylindrical volume of 6 mm diameter × 12 mm deep was used for quantification, as bone formed within the defect was of interest rather than bone formed at the periphery from stimulation during defect creation (**Supplementary Figure 14**).

### 2.6 Histological analysis of sheep study bone defects

#### 2.6.1 Processing, embedding, and sectioning of bone tissue

Samples were fixed in 10% NBF for 2-3 weeks in total to ensure complete fixation of the tissues; initially the whole distal femur, followed by the medial femoral condyle and finally the trimmed bone samples post-µCT scanning. After µCT scanning, the samples were cut in half longitudinally or into 2 – 3 pieces transversely using a water-cooled band saw (Buehler Isomet Low Speed Saw), and part used for LR white hard resin embedding and part used for wax embedding (**Supplementary Figure 15 A)**. The samples for wax embedding were cut into thinner slices, rinsed in PBS and placed in decalcification solution at RT. Decalcification was achieved using 10% ethylenediaminetetraacetic acid (EDTA) in 0.1 M Tris in dH_2_O (pH 7.3 with 5 M NaOH) for 6 weeks. The solution was changed every 3 – 4 days, with µCT scanning used to confirm complete decalcification, prior to rinsing in PBS and further processing. After dehydration through ethanol solutions (70%, 90%, 100% twice, with 30 minutes in each concentration) and chloroform:100% ethanol mix in a 1:1 ratio, followed by 100% chloroform twice, for 60 minutes in each solution, the samples were processed in a Heraeus vacutherm vacuum oven in molten paraffin wax (Histowax, Leica) at 65 °C for 1 hour. Fresh wax was added and a further 2 hours in the vacuum oven commenced. The samples were embedded in fresh molten wax and cooled at 0 – 1 °C for 2 hours before storage at 4 °C. The blocks were sectioned at 10 µm on a Microm 330 microtome (Optec, UK) (**Supplementary Figure 15 B)**. The sections were transferred via a water bath to pre-heated glass slides for 2 hours until dry and placed in the slide oven at 37 °C for 4 – 6 hours prior to storage at 4 °C.

For LR white hard resin embedding of non-decalcified bone samples, the tissue was placed in ethanol of increasing concentrations at 4°C. The bone samples were placed in 70% ethanol for 72 hours. The ethanol was changed for 80% ethanol for 24 hours followed by a further 48 hours. This was changed for 90% ethanol for 24 hours and changed to 100% ethanol for 48 hours followed by a change to fresh 100% ethanol for a further 24 hours. The tissues were placed in chloroform for 24 hours at RT to defat the tissue, and subsequently moved to 100% ethanol again for 24 hours and changed to fresh 100% ethanol for 48 hours to remove all traces of chloroform. The tissue samples were immersed in LR white resin to allow resin to surround each entire sample for 72 hours. The resin was changed for fresh resin every 3 days over two occasions (i.e., 9 days in resin) and the sample was removed from the container. The sample was placed in a 30 mL polythene container of sufficient size to allow the resin to surround the tissue to ensure complete embedding and filled to the top with LR white hard resin and accelerator (3 drops accelerator to 30 mL resin) and closed securely with the screw-on lid. The container was placed in the freezer at -20°C to cure. After 30 minutes polymerisation was complete. The samples were easily removed from the polythene container with a hacksaw (Wickes, UK) and stored at 4°C.

The resin samples were cut with a hacksaw and fine-toothed hacksaw blade, to remove excess resin and to allow the samples to fit into the holder for the band saw (Buehler Isomet Low Speed Saw) (**Supplementary Figure 15 C**). Thin ∼1 mm wide sections were cut on speed setting 10 and glued to Perspex (3 inch long × 1 inch wide × 3 mm deep) rectangles (to produce non-shatter slides) using Loctite^®^ superglue (Henkel). The plastic backing was kept on the Perspex slide on the opposite side from the sample to prevent scratches. The glue was allowed to dry for at least 24 hours before the slide was turned over and a blob of Blu Tack^®^ (Bostik) used at either end of the slide to allow the slide to be held, while keeping fingers away from the rotating plate, while grinding the resin slice. The slide was placed on a grinder/polisher machine (MetPrep 20 DVT) at 225 rpm using sandpaper of 1200 and 4000 grades to grind the sample to ∼100 µM (**Supplementary Figure 15 D**). Staining methodology was as described for wax without the prior Histoclear, rehydration, or dehydration steps.

#### 2.6.2 Histology staining of tissue samples

##### 2.6.2.1 Alcian blue/Sirius red

The method has been previously described (14). In brief, the wax embedded tissue sections on slides were rehydrated through Histoclear (2 × 7 minutes), ethanol solutions of 100% (twice) to 90% to 50% (2 minutes each), followed by immersion in water. Weigert’s Haematoxylin was applied for 10 minutes, removed by immersion 3 times in acid/alcohol (5% HCl/70% ethanol) followed by washing in the water bath for 5 minutes. Slides were immersed in 0.5% Alcian blue 8GX in 1% acetic acid for 10 minutes, 1% molybdophosphoric acid for 10 minutes, followed by rinsing with water prior to staining with 1% Picrosirius Red (Sirius Red) for 1 hour. Excess stain was rinsed off in water and the slides were dehydrated with increasing concentrations of ethanol of 50%, 90%, 100% twice and Histoclear twice for 30 seconds in each prior to mounting with dibutyl phthalate xylene (DPX) and a glass coverslip and allowed to dry.

##### 2.6.2.2 Goldner’s Trichrome

The method has been previously described (14). In brief, the method followed is identical to Alcian blue/Sirius Red staining to the point of acid/alcohol immersion and washing in water for 5 minutes. Ponceau Acid Fuchsin/Azophloxin (Sigma) was applied for 5 minutes followed by a 15 second wash in 1% acetic acid. Phosphomolybdic acid/Orange G was applied for 20 minutes followed by another 15 second wash in 1% acetic acid. Light Green was applied for 5 minutes followed by the third 15 second wash in 1% acetic acid. The sections on the slides were dehydrated in ethanol 90% and 100% twice and Histoclear twice for 30 seconds each prior to mounting with DPX and a glass coverslip and allowed to dry.

##### 2.6.2.3 Alizarin red/light green staining

The Perspex slides with resin embedded samples were placed on a slide rack and stained by application of Alizarin red S (40 mM) stain solution for 2 minutes. The slides were blotted and air-dried.

##### 2.6.2.4 Tartrate-resistant acid phosphatase (TRAP) staining

The method has been previously described (17). For the visualisation of osteoclast activity, TRAP Basic Incubation Medium (9.2 g Sodium Acetate Anhydrous, 11.4 g L-tartaric Acid, 950 mL dH_2_O, 2.8 mL glacial acetic acid, pH adjusted with 5 M sodium hydroxide to pH 4.7-5.0 and made up to 1 L with dH_2_O and Naphthol AS-MX Phosphate substrate (2 mg Naphthol AS-MX Phosphate to 0.1 mL of ethylene glycol monoethyl ether) were prepared. The final staining solution was prepared from 20 mL of TRAP basic incubation medium, 0.1 mL of Naphthol AS-MX Phosphate substrate and 12 mg of Fast Red Violet LB Salt, prewarmed to 37°C in a drying oven. The slides were dewaxed and rehydrated prior to staining with TRAP staining solution applied to each slide and incubated at 37°C for 30 minutes, checking red positive stain development on positive control slides. The slides were rinsed in dH_2_O and counterstained with 0.02% Fast Green stain for 30 seconds, rinsed in dH_2_O, dehydrated through 50%, 90%, 100% (twice) and Histoclear (twice) for 30 seconds in each solution. Slides were mounted with DPX and a glass cover slip applied and allowed to dry.

#### 2.6.3 Image acquisition of histology slides

Histological wax embedded samples were imaged using the Zeiss Axiovert 200 digital imaging system using bright field microscopy with the halogen bulb. Images were captured using the Axiovision 4.2 imaging software. The Olympus Virtual Slide System VS 110 was used to image the slides and images captured using the Olympus VS-Desktop 2.9.1 software. The resin embedded samples were imaged using Leica TCS-SP8 microscope using dual excitation at 405 nm and 458 nm and primary detection of 480-560 nm and red was set at 600-690 nm for fluorescence detection. The resin embedded samples stained with alizarin red were imaged using an Epson Perfection V700/V750 scanner.

#### 2.6.4 Statistical analysis

Analysis and graphical presentation were performed using GraphPad Prism 9, version 9.2.0. P values <0.05 were considered significant. Graphical representation of significance as follows: ns is no significant difference, *p<0.05, **p<0.01, ***p<0.001, ****p<0.0001. All data presented as mean and standard deviation (S.D.).

## 3. Results of the ovine femur defect study

### 3.1 Sheep health and welfare parameters

#### 3.1.1 The sheep remained healthy during the 13 week femoral condyle defect study

Sixteen sheep were used in the study. One sheep died <24 hours postoperatively of an unknown cause, the remaining fifteen sheep displayed excellent recovery. No infections were observed and animals received analgesia and antibiotics as planned. One sheep (empty defect and an autograft-filled defect) became lame post IM injection of tetracycline, due to suspected muscle pain, which resolved with oral administration of meloxicam (Metacam^®^, Boehringer Ingelheim). Weight and body condition scores were recorded for each animal prior to surgery and at the week 13 end point for monitoring animal health during the study. Body weights of all animals increased to an average weight of 85.0 ± 7.7 kg over the 13 week study. All animals had static or increased body condition scores at time of sacrifice of average 3.20 ± 0.37. This suggested that no issues were encountered with regards to the safety of the surgical procedure or the materials implanted.

### 3.2 Analysis of bone formation and biocompatibility of materials within the femoral defect sites

#### 3.2.1 Bioactive BMP-2 and Laponite coated PCL-TMA900 scaffolds produced significantly more bone around the Laponite/BMP-2 coated PCL-TMA900 scaffolds than the uncoated PCL-TMA900 scaffolds

The study demonstrated that the empty defects did not fill with bone to fully repair the defect, rather, irregular, sparse, bone formation from the base, top or sides of the defect was observed which would not be conducive to bridging of the defect. The autograft implants within the defect formed a dense mass of bone tissue. The uncoated PCL-TMA900 scaffolds displayed minimal bone formation, with the scaffold inhibiting bone formation within the defect, evidenced by the sparse bone spicules entering the scaffold and absence of new bone from the base or top of the defect. In comparison, bone formation was observed around the Laponite/BMP-2 coated scaffolds with bone forming over the shape and pore channels of the scaffold. This was observed to be consistent in 50% of the implants, with less bone formed at one end of the implant closest to the periosteal side of the defect in the other 50% of sheep (**Figure 1 A**).

**Figure 1:**
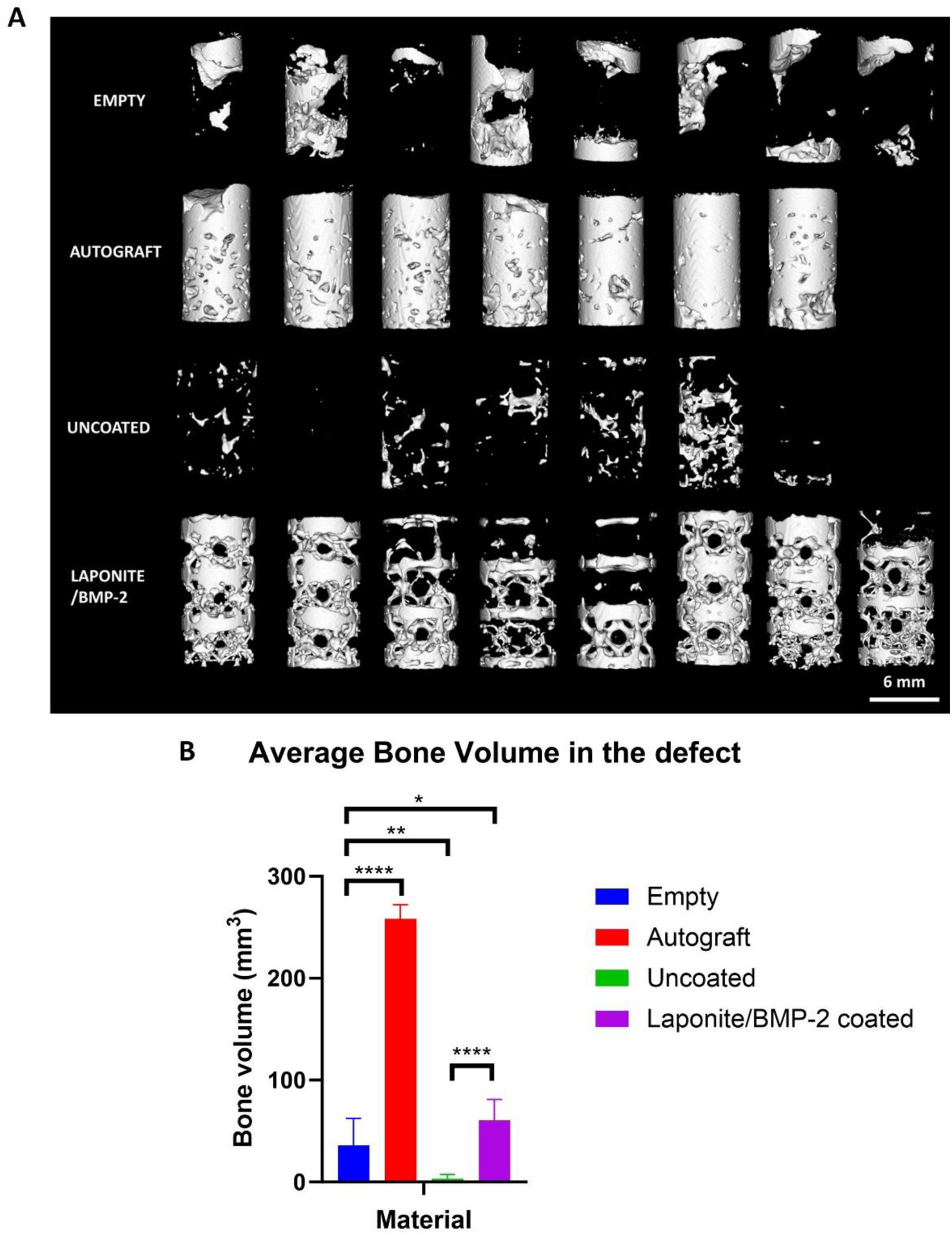
Images of representative samples and quantification of bone volume using µCT (A) Reconstructed images at 40 µm showing bone formed within the 6 mm x 12 mm volume within the centre of the bone defect in each group. The empty defects displayed bone formation, in regions, especially from the top or bottom of the defect with a large void still present. The autograft group showed dense bone, while the uncoated scaffold showed discrete spicules of bone within the scaffold, which was not consistent between sheep. The Laponite/BMP-2 coated scaffolds showed bone formation on the surface of the scaffold, allowing the shape of the scaffold and pores to be determined. Scale bar 6 mm. (B) The average bone formed within the defect region quantified was significantly greater in the autograft control (n = 7) and Laponite/BMP-2 coated (n = 8) groups, while the uncoated scaffold (n = 7) showed significantly less bone formation than when the defect remained empty (n = 8). One-way ANOVA with Dunnett’s multiple comparisons test, mean and S.D. shown, *p<0.05, **p<0.01, ****p<0.0001. Analysis determined there was a statistically significant difference in bone formation between the Laponite/BMP-2 coated (n = 8) and uncoated (n = 7) PCL-TMA900 scaffolds. One-way ANOVA with Šidáks multiple comparisons test, mean and S.D. shown, ****p<0.0001.

The bone formation response to the autograft, empty defect, uncoated scaffolds and Laponite/BMP-2 coated scaffolds were quantified. The volume of bone which could theoretically fill the selected defect volume (i.e., 6 mm × 12 mm defect) was calculated at 339.12 mm^3^. The scaffold material itself had a volume of 185.08 mm^3^ leaving 154.04 mm^3^ available for infill of new bone in the 6 mm diameter × 12 mm deep cylinder. At 40 µm reconstruction parameter, in comparison to the empty defect with a mean bone volume of 35.88 mm^3^, the autograft scanned *ex vivo* showed significant mean bone volume of 258.24 mm^3^ unsurprisingly, as the defect was filled with impacted bone graft. The uncoated PCL-TMA900 scaffolds displayed significantly lower net bone growth than the empty defect, presumably due to the scaffold limiting bone infiltration/formation, while providing mechanical support, with a mean bone volume of 3.08 mm^3^. In marked contrast, the Laponite/BMP-2 coated PCL-TMA900 scaffolds displayed significant bone formation with a mean volume of 60.75 mm^3^ compared to the empty defect (**Figure 1 B**). This equates to 39.4% of the available volume outside the scaffold being filled with bone tissue in the Laponite/BMP-2 coated scaffold group. On comparison of the uncoated and Laponite/BMP-2 coated scaffolds, there was a significant difference in bone volume formed due to the coating on the scaffolds (**Figure 1 B**).

When reconstructions were performed at 20 µm the bone formed was similar to that viewed at 40 µm, with bone in sporadic regions of the empty defect and filling the autograft-filled controls. The uncoated scaffold evidenced spicules of bone, which were irregular and inconsistent, whereas the Laponite/BMP-2 coated scaffolds showed the same pattern of bone formation as at 40 µm, indicating that the bone formed was of significant quantity to be seen clearly at both resolutions, with bone forming an outer shell over the scaffold shape (**Figure 2 A**). Upon quantification, the Laponite/BMP-2 coated PCL-TMA900 scaffolds displayed bone formation with a mean volume of 52.48 mm^3^, however, this was not found to be a significant difference in quantity compared to the empty defect with a mean of 34.74 mm^3^. The uncoated scaffold showed significantly less bone formation equating to a mean volume of 2.68 mm^3^, while the autograft had a mean volume of 224.36 mm^3^ and therefore showed a significant difference in bone volume in the defect, than in the empty defect, in agreement with 40 µm resolution findings (**Figure 2 B**). In agreement with analysis at 40 µm, there was a significant difference in bone volume within the defect between the uncoated and Laponite/BMP-2 coated scaffolds (**Figure 2 B**).

**Figure 2:**
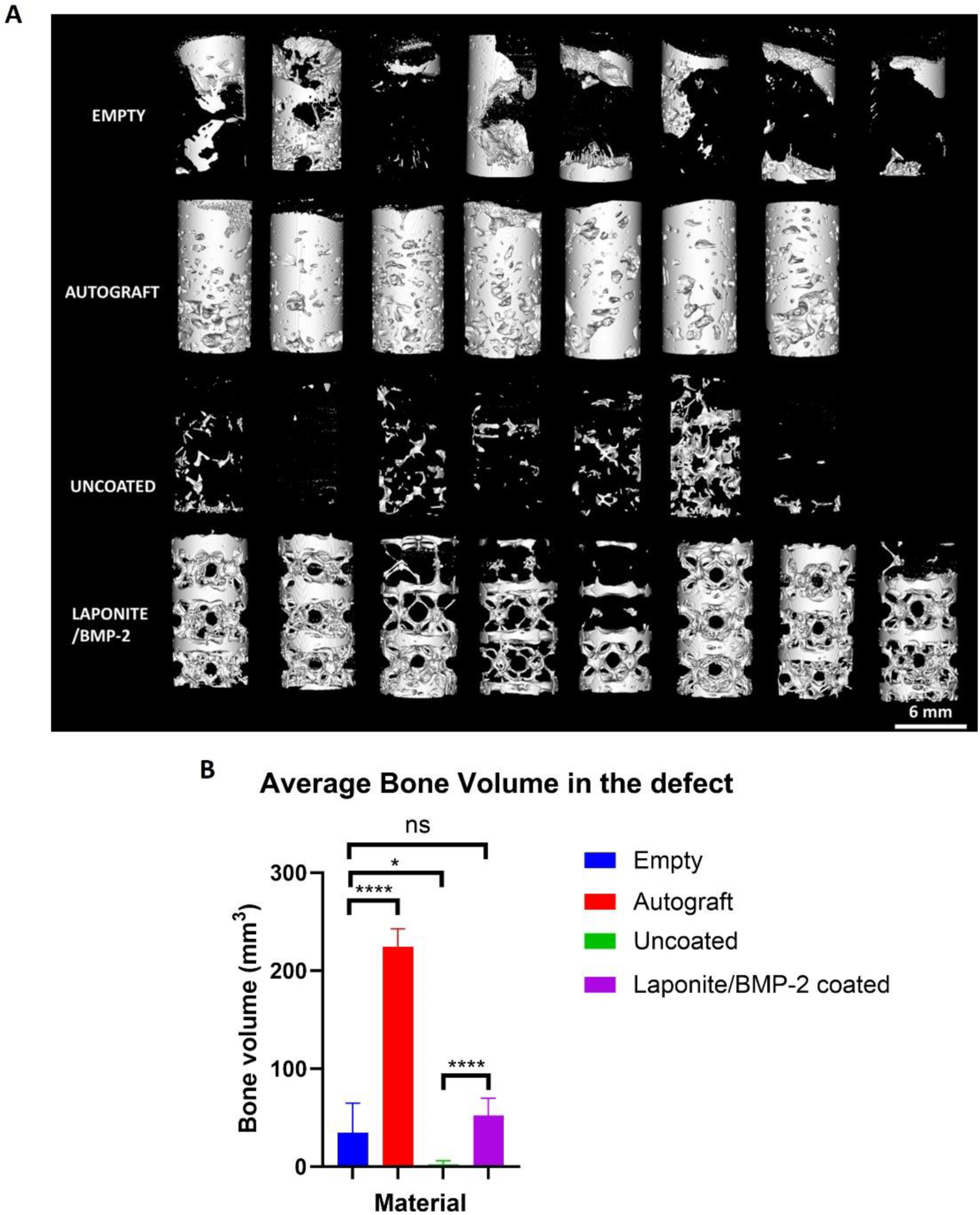
Images of representative samples and quantification of bone volume using µCT (A) Reconstructed images at 20 µm showing bone formed within the 6 mm x 12 mm volume within the centre of the bone defect in each group. The images were comparable to images obtained at 40 µm, with the bone on the coated scaffolds formed at the site of the Laponite/BMP-2 coating creating a pattern and spanning the defect length in 50% of the implants. Scale bar 6 mm. (B) The average bone formed within the defect region quantified was significantly greater in the autograft control (n = 7), while the uncoated scaffold (n = 7) showed significantly less bone formation than when the defect remained empty (n = 8). Due to the higher resolution the Laponite/BMP-2 coated (n = 8) group did not display a significant difference in bone volume to the empty defect, however, the µCT images show a defined, scaffold-templated pattern of bone distribution compared to the empty defects. One-way ANOVA with Dunnett’s multiple comparisons test, mean and S.D. shown, ns; non-significant, *p<0.05, ****p<0.0001. There was again a significant difference in bone formation between the Laponite/BMP-2 coated (n = 8) scaffolds and the uncoated (n = 7) PCL-TMA900 scaffolds with greater bone formation when the coating was applied. One-way ANOVA with Šidáks multiple comparisons test, mean and S.D. shown, ****p<0.0001.

#### 3.2.2 Gross analysis revealed no complications with implantation of the PCL-TMA900 material or the Laponite/BMP-2 coating

The bone samples were transected longitudinally or transversely into sections. The bone around each defect appeared healthy with no evidence of haemorrhage or infection. The empty defects contained granulation tissue within the defect with regions of bone at the edges. The autograft had formed dense bone within the defect with the outline of the defect clearly visible. The uncoated scaffold contained regions of soft, granulation-type tissue within central regions of the scaffold and bone around the periphery where the bone had integrated with the scaffold. The Laponite/BMP-2 coated scaffolds contained bone within the scaffold between the complex structure of the octet-truss PCL-TMA900 struts (**Figure 3**).

**Figure 3:**
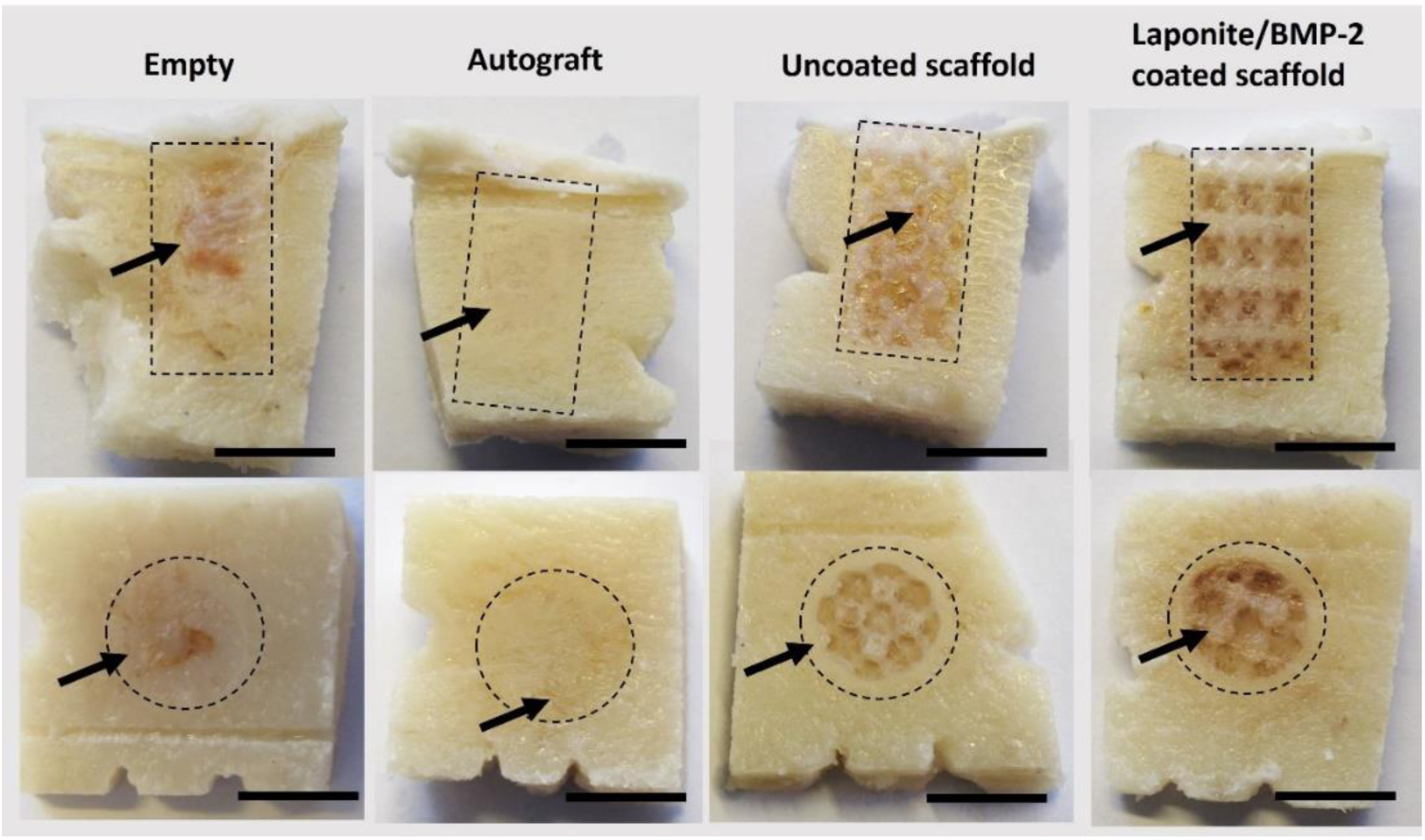
Representative images of tissue formed within the defect site in each group in longitudinal and transverse sections. The defect region is shown by the black dotted cylinder/circle in each image. The empty defect contained granulation tissue filling most of the defect, while the autograft healed as a region of dense bone (black arrows). The uncoated scaffold material can be seen with granulation tissue within the defect (top image black arrow) and a dense ring of bone surrounding the scaffold (bottom image black arrow). The Laponite/BMP-2 coated scaffold showed parallel regions of bone formed between the struts of the scaffold (top image black arrow) and around the scaffold shape shown in the transverse section (bottom image black arrow). Scale bar 8 mm.

### 3.3 Histological analysis of the tissue formed in each defect group

#### 3.3.1 Staining of tissues and fluorescence microscopy confirmed the µCT findings with obvious bone formation in the Laponite/BMP-2 coated scaffold group compared to the uncoated scaffold group

Histological analysis using alizarin red staining highlighted the bone formation within each group, confirming the observed µCT results (**Figure 4**). In the empty defect, bone was observed emerging from the base of the defect. Dense bone was observed in the autograft group with almost no bone in the uncoated scaffold and bone seen bridging the defect in the Laponite/BMP-2 coated group in the example sections shown. Tetracycline labelling was employed to examine the newly formed bone (fluorescent green), which can be seen in each group. It proved difficult to accurately determine two distinct tetracycline label lines formed at 6 and 11 weeks to determine the bone formation rate although, this was deemed less important given the clear evidence of bone formation on µCT analysis with the new fluorescent bone indicative of defect healing. The empty group and uncoated PCL-TMA900 groups displayed new bone at the periphery of the defect where dense bone was forming. The autograft group contained regions of fluorescence within the implanted bone and the Laponite/BMP-2 coated scaffold contained fluorescent bone within the pores of the scaffold and new bone clearly evidenced around the perimeter of the scaffold.

**Figure 4:**
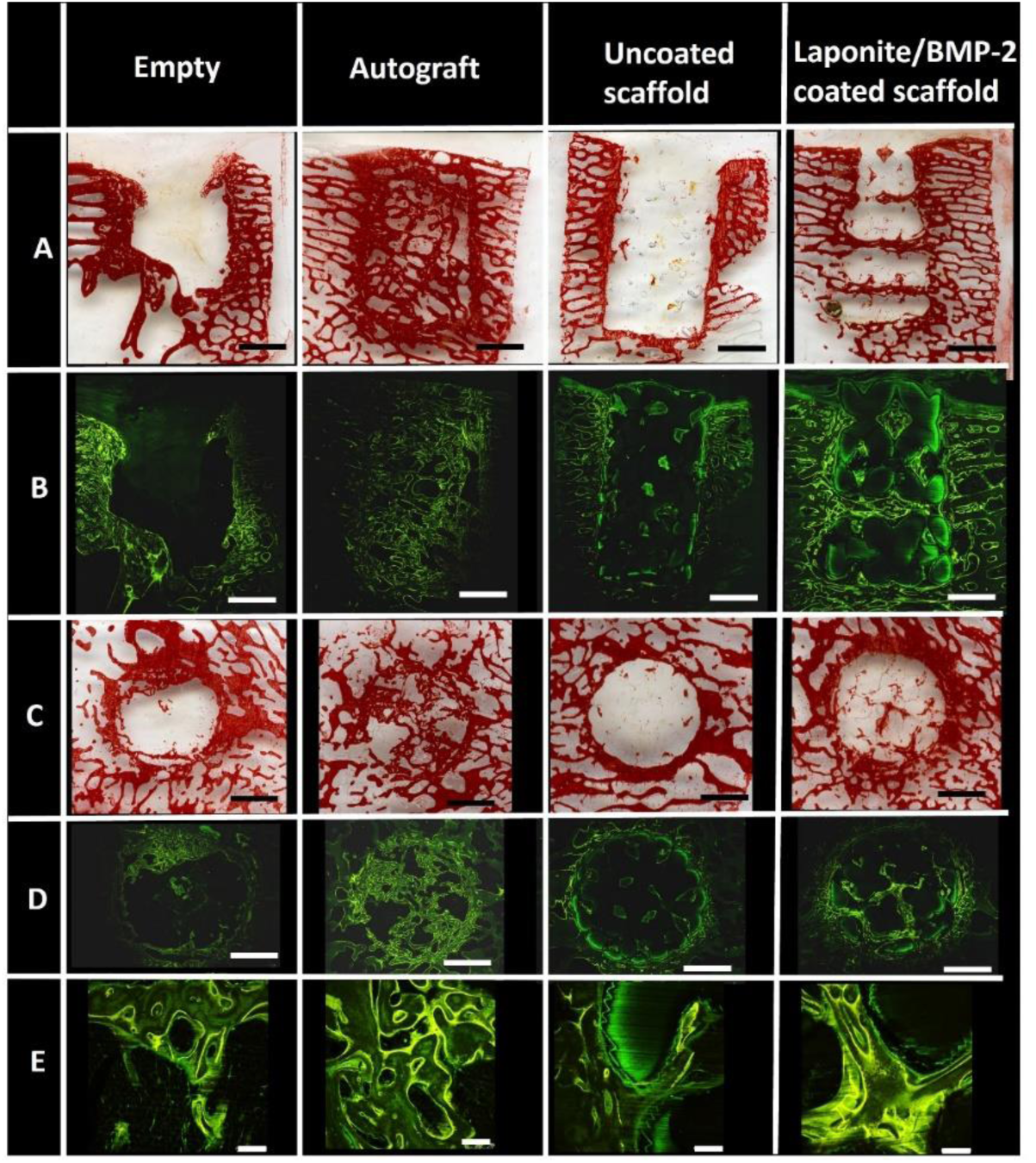
Representative images of histology of resin embedded tissue from each group. Row A: Alizarin red staining of the defect groups with the bone formed within the defect shown in red. Row B; Fluorescence images of bone sections showing the new bone (fluorescent green) forming at the periphery and within the defects. Row C; Alizarin red staining of transverse sections showing bone at the edges of the empty defect and uncoated scaffold, while there is a network of new bone in the autograft control and a criss-cross shape of bone formed in the Laponite/BMP-2 coated scaffold group. Row D; Fluorescence images of fluorescent green new bone, mirroring the bone seen within the defects in row C. Row E; Magnified images of row D showing new bone at the edge of the empty defect, dense implanted bone in the autograft group, new bone at the periphery of the uncoated scaffold group with the scaffold material seen as jagged lines and new bone formed at the centre of the defect in a cross-shape in the Laponite/BMP-2 coated scaffold group. Rows A-D scale bar 2 mm, Row E scale bar 250 µm.

#### 3.3.2 Laponite/BMP-2 coated scaffolds contain predominantly bone and marrow tissue, while the uncoated scaffolds contained mainly fibrous tissue

Soft tissue composed of collagen fibres and marrow tissue with discrete regions of bone formation around the periphery was found within the empty group, while dense bone with almost no marrow cavities in places was observed in the autograft control group. The uncoated PCL-TMA900 scaffolds displayed a rim of bone around the scaffold periphery, with predominantly fibrous tissue extending into the scaffold construct observed as well demarcated ‘fingers’ of tissue and fibrous tissue between the struts of the scaffold. In contrast, the Laponite/BMP-2 coated group showed bone and marrow tissue extending across the scaffold and around the polymer struts of the scaffold (**Figure 5**). Further images are shown in **Supplementary Figures 16, 17 and 18**.

**Figure 5:**
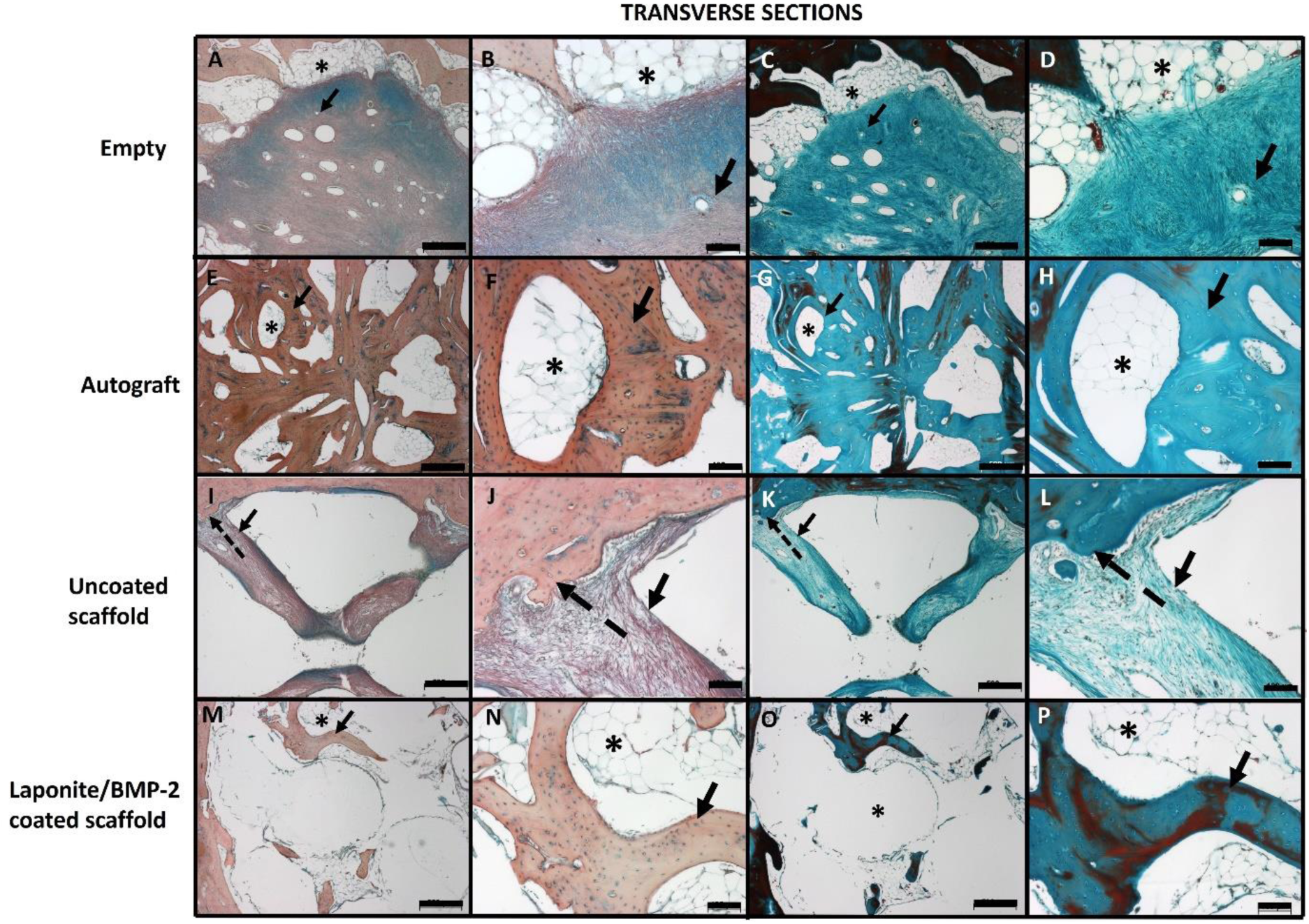
Representative images of histology of transverse sections of defects from each group. (A, B, E, F, I, J, M, N) Alcian blue and Sirius red staining (C, D, G, H, K, L, O, P) Goldner’s trichrome staining. A–-D; empty defect with pink/green collagen fibres (black arrows) and bone and marrow tissue surrounding the defect (asterix). E–-H; Autograft with irregularly aligned bone struts (black arrows) within the defect and marrow cavities (asterix). I–-L; uncoated scaffold with bone around the scaffold (dashed arrows) and collagen fibres (black arrows) extending into the scaffold with a clear demarcation between bone and fibrous tissue. M – P; Laponite/BMP-2 coated scaffold with bone tissue (black arrows) extending from the periphery into pores of the scaffold with surrounding marrow tissue (asterix). A, C, E, G, I, K, M, O scale bar 500 µm and B, D, F, H, J, L, N, P 100 µm.

TRAP staining showed osteoclast activity in the dense bone surrounding the empty defect in discrete cavities, while the autograft sample had no osteoclast activity detected. The uncoated scaffold showed osteoclast activity at the junction between the peripheral bone and the fibrous tissue extending into the scaffold, compared to minor osteoclast activity noted within the pores of the Laponite/BMP-2 coated scaffold. This confirmed the defect site was undergoing minimal remodelling in each group at the end point of the study (**Figure 6**).

**Figure 6:**
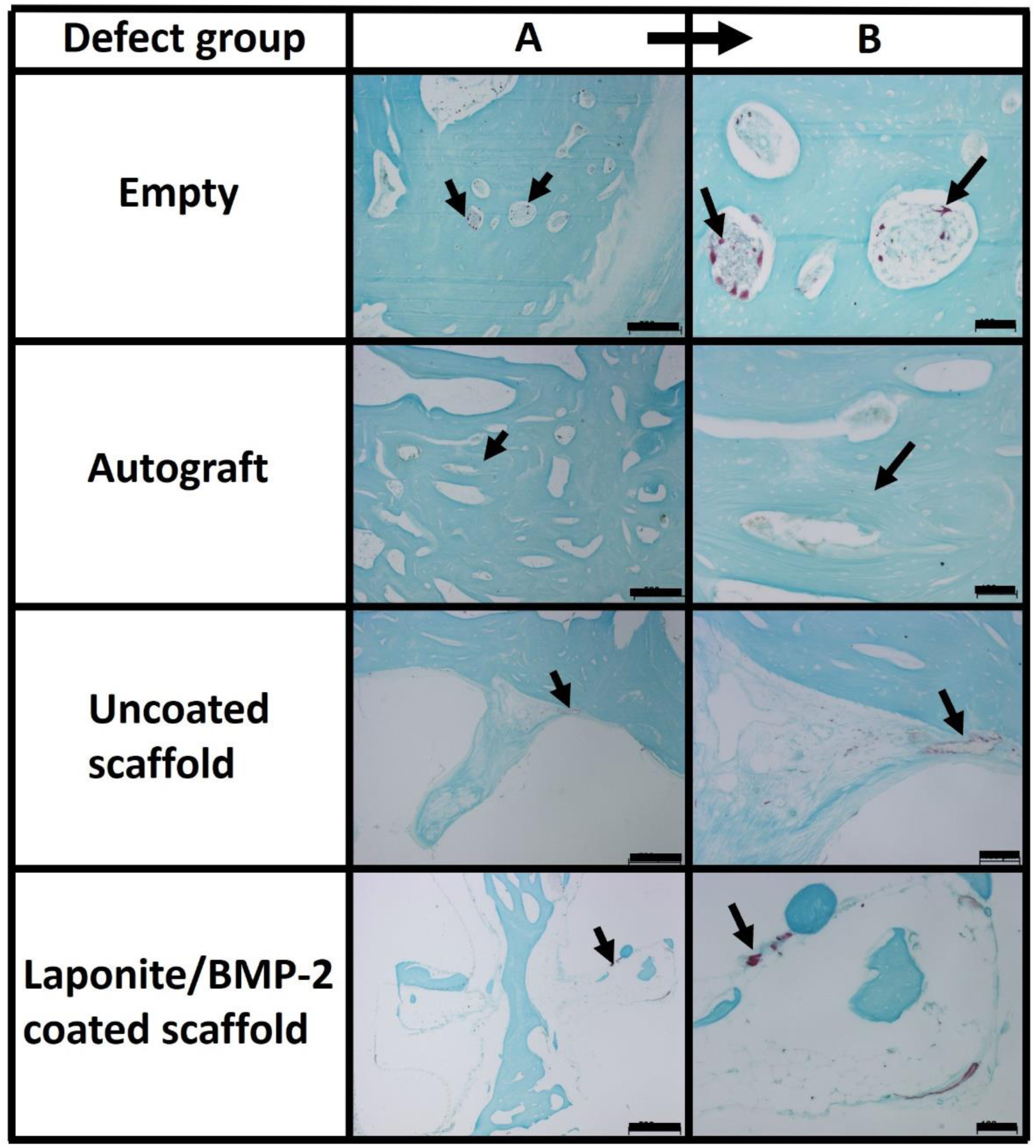
Representative images of TRAP staining of sections to visualise osteoclasts in the bone remodelling process. In the empty defect group, osteoclasts were visualised in the bone surrounding the defect (black arrows) but not within the defect itself. The autograft group showed no osteoclasts within or surrounding the defect site, with osteocytes (black arrow) seen in a regular pattern around marrow cavities in the defect. The uncoated scaffold had osteoclasts at the junction between bone and fibrous tissue (black arrows), but not within the scaffold region. The Laponite/BMP-2 coated scaffold contained osteoclasts at the scaffold surface between marrow and bone formed in response to the coating. (A) scale bar 500 µm (B) scale bar 100 µm.

## 4. Discussion

In the current study, we hypothesised that Laponite and BMP-2 coated PCL-TMA900 scaffolds would produce significantly more bone than the uncoated PCL-TMA900 scaffolds, with the coated scaffolds enhancing healing of a critical-size bone defect. As shown in our previous study, Laponite nanoclay adhered to the PCL-TMA900 material and subsequent immersion of the Laponite coated scaffolds in BMP-2 solution enabled sequestration of the BMP-2 by the Laponite coating to deliver active BMP-2. Studies confirmed the ability for Laponite to remain adhered to the PCL-TMA900 scaffold for implantation in sheep, while ELISA results confirmed BMP-2 binding to the nanoclay coated scaffold to a saturation point; likely linked to the surface area of the scaffold and mass of adherent Laponite. This produced a bioactive, osteoinductive scaffold of macro size able to be scaled up for clinical evaluation. In contrast to the coated scaffold, the uncoated scaffold did not facilitate bone healing, while providing some mechanical support to the surrounding bone.

In another study by Berner and colleagues, PCL-tricalcium phosphate scaffolds were tested with allogenic mesenchymal progenitor cells, orofacial osteoblasts or axial skeleton osteoblasts to assess bone regeneration in a sheep tibial defect model, showing safety and efficacy of the implanted cells and scaffold material (31). A key point of the current study is the use of an acellular scaffold material to bypass the necessity of cell culture and the associated barriers with the use of cells for tissue regeneration, instead harnessing the regenerative potential of the *in situ* skeletal cell populations to respond to the osteogenic scaffold itself. The PCL-TMA900 used in this study has been tested using *in vitro* cytocompatibility assays, the chorioallantoic membrane (CAM) assay to test biocompatibility, in murine subcutaneous implantation studies and murine femur defect studies prior to evaluation in this ovine femur defect study (14, 17). The compatibility observed with the ovine bone tissue, grossly and histologically, is therefore unsurprising given the previous supportive avian and rodent *in vivo* studies. Furthermore, our recent studies confirmed significant bone augmentation *in vivo* using mouse subcutaneous and segmental defect models using the Laponite/BMP-2 coating on the scaffold, facilitating scale-up in this ovine bone defect model (14, 17). The fold increase in defect size/cylinder volume from the murine femur defect to sheep defect size was five times, and the sheep scaffold was over ten times the surface area of the mouse femur scaffolds (113.9 mm^2^) detailed previously (17). In the literature, Stro-4+ ovine BMSCs with a bovine ECM/PCL hydrogel scaffold were tested *in vitro* and on the CAM prior to an *in vivo* tibial defect study in sheep, however, the promising results of the preliminary testing did not translate to bone formation in the sheep model (32). This highlights the need for robust testing prior to scale-up with clear evidence of efficacy, as *in vitro* and *in vivo* findings do not always correlate (33). The use of 3D printing to manufacture an osteoconductive calcium phosphate scaffold illustrated 3D printed scaffold use in sheep as a model for human critical size defects (22).

Upon quantification, the Laponite/BMP-2 coated PCL-TMA900 scaffolds displayed bone formation, with a significant difference compared to an empty defect when analysed at 40 µm, however, this was not found to be a significant difference compared to the empty defect in bone quantity when analysed at 20 µm. It is important to note that the scaffold when used clinically, would enable the new bone to span the defect compared to the irregular islands of bone formed in the empty defect, thereby restoring some of the load-bearing capacity of the bone. There was a consistent, clear, significant difference between the uncoated scaffold and Laponite/BMP-2 coated scaffold use regarding the bone volume seen grossly and following examination using µCT, with histology confirming the tissue type formed within the defect, i.e., fibrous versus bone tissue. The histology revealed fibrous tissue extending from the bone surrounding the uncoated scaffold, while the Laponite/BMP-2 coated scaffold showed bone formation throughout the scaffold surface, as a consequence of the osteogenic nature of the Laponite/BMP-2 coating. The results confirm that while skeletal stem/progenitor cells are at the site of injury, a scaffold alone would not be sufficient to form bone tissue, at least not in a mechanically stable site. The autograft formed dense tissue with small regions of marrow tissue, but this would be mechanically distinct to native trabecular bone due to the density of trabecular bone within the defect site. The empty defect was filled with granulation-type tissue of collagen fibres, with marrow tissue towards the base of the defect, hence clinical human large bone defects are not left unfilled due to this sporadic, unreliable bone formation. TRAP staining to assess the remodelling process due to osteoclast activity revealed minor, sporadic, activity in the dense bone surrounding the empty defect and uncoated scaffold and within the pores of the Laponite/BMP-2 coated scaffold. However, no obvious remodelling was noted on sections of the autograft-implanted defect. This may be due to lack of mechanical stimulus at the femur condyle site; as when scaffolds were implanted into mouse femur defects, the instability and forces of locomotion are thought to have stimulated osteoclastic remodelling (17).

The use of a large animal model allows evaluation and assessment of the biomechanics, degradation, growth factor release and pharmacokinetics at play to be assessed, with sheep having similar body weight and bone turnover characteristics to humans (34). Sheep of 3-4 years old have rapidly forming, plexiform bone which is a combination of woven and lamellar bone and only show haversian remodelling at 7-9 years of age, firstly at the caudal aspect of the femur and the diaphysis of the radius and humerus, making older sheep bone more similar to humans (35). Skeletally mature sheep were used in this project as it had been shown previously that the empty defect does not heal itself over the 13 week duration of the study (30). In the current study, the empty defect filled with fibrous tissue and marrow tissue at the edges and bottom of the defect, with no mature bone seen within the centre of the defect or spanning the height of the defect. The definition of a critical-sized defect varies in the literature, as the smallest defect to not permit spontaneous healing is often not tested against other defect sizes in the exact same model/species/site and the end point of the study often is before the natural end of the animal’s life (36). The sacrifice time point selected in the current study was 13 weeks, as it was expected that significant bone growth should have occurred due to the rate of bone remodelling.

This study used a critical-sized defect to enable examination of the effect of the scaffold and BMP-2 growth factor on bone healing without fracture fixation or differences in stability between sheep, which may be seen following plating or external skeletal fixator application for large segmental critical-sized defects. With this model (large segmental defect), sheep are reported to have several challenges including post-operative fractures due to stress fractures developing from drill holes and due to brittle bones, difficulty in using the optimal screw diameter due to 3.5 mm screws being the correct size but inadequate strength, and postoperative issues such as over-exertion, getting up and down or with the environment (37). The use of a sling to prevent the animal trying to fully weight bear on the operated limb to mimic the clinical scenario for humans postoperatively has been developed. The sling allows the amount of weight borne by the limb to be adjusted and then kept constant, whilst maintaining comfort and to avoid detriment to normal bodily functions and stress and improve welfare (38). Pain scales for sheep have been published and we used grimace and locomotion scoring, alongside body condition scoring and weight measurements to ensure the health and welfare of the sheep prior to and during the study (39). Due to the defect being non-articular and the limb stable without fixation, the sheep appeared to be comfortable without the need for interventions to support the limb or aid weightbearing, allowing bilateral defects to be made as the sheep were fully-weight bearing and freely ambulatory post-operatively.

This metaphyseal bone location allowed the scaffold to be within cancellous bone and not contact the intramedullary marrow which would be a rich source of skeletal stem cells (40). A model allowing assessment of cancellous and cortical bone response to material has been developed, with 16 defects possible within the one animal, with defects on one side made followed by the other side at a later point to allow different time point evaluation. The use of the epiphysis of the femur and humerus to create 6 mm diameter, 15 mm deep defects and unicortical metacarpal and metatarsal defects comprising the cortical defects of 6 mm diameter are made, with refinements made when making the defects to reduce the risk of fracture formation (41). This model would allow more data to be collected per sheep and potential determination of the optimal implantation site (i.e., cortical and/or cancellous bone) or capabilities of a novel scaffold material. However, the metaphysis is a stable, relatively safe site of bone defect creation. Therefore, the metaphysis was used in our study to reduce the risk of complications and confounding factors, such as the differences in weight bearing between forelimbs and hindlimbs.

The use of haematological markers of bone turnover e.g. alkaline phosphatase and minerals have been studied as a non-invasive method of monitoring bone turnover in research settings (42). The use of ultrasound was found to be an adjunct to the use of CT when monitoring tibia bone healing, but ultrasound was found to overestimate the mass of new bone compared to the CT findings (43). In this current study, bone formation following tetracycline administration at weeks 6 and 11, was examined however, the 3D geometry of the bone when resin embedded and potential for possible misalignment of the bone in the resin, can preclude with any certainty that two distinct lines of tetracycline have formed. Nevertheless, in the current study, the tetracycline labelling was found to label the new bone, with clear differences between the 4 groups studied which mirrored the alizarin red staining for bone tissue.

The PCL-TMA900 material is known to be hydrophobic with limited cellular attachment, while the degradation profile of the material will be important for clinical translation. There appeared to be minimal macroscopic degradation in this sheep study, as it is known that PCL can take months to years to degrade (44). It would be interesting to implant the PCL-TMA900 scaffold within an ovine tibial defect model to subject the scaffold to greater forces and to study the degradation over a longer timeframe. The Laponite/BMP-2 coating induced bone to form over the surface of the scaffold, however, this appears to become static once this bone has formed. It is interesting to speculate, degradation of the PCL-TMA900 material may allow the micro-instability to signal to the resident osteocytes and osteoblasts within the ossified layer on the scaffold to produce further bone to stabilise the defect. Therefore, a positive feedback cycle may ensue leading to complete healing of the bone defect, incorporating or replacing the PCL-TMA900 scaffold material over time. Long term assessment of results can be difficult due to practical and financial considerations, however, long term studies of up to 12 months have been reported in bone tissue engineering to determine the efficacy of the scaffold, bone quality produced and degradation of the material (45). The degradation of a material and macrophage response has gained more importance in directing the macrophages towards an M2 pro-regenerative phenotype, rather than an inflammatory M1 type. Wu and colleagues reported if a material degrades too quickly, macrophage polarisation can be disrupted and bone healing abated (44).

Further work will address refinement of the Laponite coating process for a variety of materials using different concentrations or methods of application. It is clear such an approach could be applied to other growth factors such as vascular endothelial growth factor to enhance vascularisation in non-healing sites. The inactivation of BMP-2 when left adhered to the Laponite for a period (i.e., 24 hours), as evidenced by previous *in vitro* and mouse subcutaneous implantation study work, requires further investigation and optimisation especially from a translational perspective (14). Advancing this work, the PCL-TMA900 material and Laponite/BMP-2 coating warrant further examination through ISO testing to translate the material scaffold and Laponite/BMP-2 coating towards clinical application in the repair of large bone defects.

## 5. Conclusion

This study investigated the use of a ready-coated, osteogenic, acellular scaffold within a critical-sized ovine metaphyseal bone defect model to confirm the biocompatibility and efficacy of bioactive Laponite/BMP-2 on rigid PCL-TMA900 polymer scaffolds for bone repair at scale. The Laponite/BMP-2 coating induced bone formation on the PCL-TMA900 scaffold within the defect, as determined by µCT imaging and histology compared to uncoated PCL-TMA900 scaffolds. The current studies confirm the efficacy of a biomimetic osteoinductive octet-truss scaffold evaluated in a large animal critical sized defect model and offers exciting potential for future translational evaluation and patient application for an increasing aging demographic.

### Credit author statement

Callens, S. J. P.: designed the scaffold shape, Wojciechowski, J. P. and Echalier, C. prepared and printed the PCL-TMA900 scaffolds, Marshall K. M. conducted the *in vitro* experiments, the ovine femur defect study, data collection and analysis, histology, and wrote the paper. M^c^Laren, J. organised, performed and supervised the ovine femur defect study and edited the paper. Dawson J. I supplied the Laponite and edited the paper. Kanczler J. M., Rose, F. R. A. J., Stevens, M. M., Dawson J. I., and Oreffo R. O. C. conceptualisation, funding acquisition, study supervision and editing of the manuscript.

### Data and materials availability

All data associated with this study are presented in the paper or the Supplementary Materials. All raw data is available upon request.

## Acknowledgements

Research support for this study from the Biotechnology and Biological Sciences Research Council (BBSRC BB/P017711/1), the UK Regenerative Medicine Platform Acellular / Smart Materials – 3D Architecture (MR/R015651/1) and University of Southampton is gratefully acknowledged as well as useful discussions with current members of the Bone and Joint Research Group in Southampton, UK. Mick Baker and the staff of the Biomedical Services Unit at the University of Nottingham supported all animal work. We thank Professor Nicholas Evans (University of Southampton) and Professor Manuel Salmeron-Sanchez (University of Glasgow) for helpful discussions on the programme of work. Dr Vineetha Jayawarna (University of Glasgow) and the University of Glasgow Veterinary School are acknowledged for organising the EO sterilisation of the PCL-TMA900 scaffolds. Dr Akemi Nogiwa Valdez is acknowledged for proof reading of the manuscript. For the purpose of open access, the author has applied a ‘Creative Commons Attribution’ (CC BY) license to any Author Accepted Manuscript version arising.

Appendix A. Supplementary data

Supplementary data to this article can be found online.

