## Supplemental information for "Bioactive coatings on 3D printed scaffolds for bone regeneration: Use of Laponite^®^ to deliver BMP-2 in an ovine femoral condyle defect model"

### Supplementary information

#### The scaffold shape, porosity and dimensions

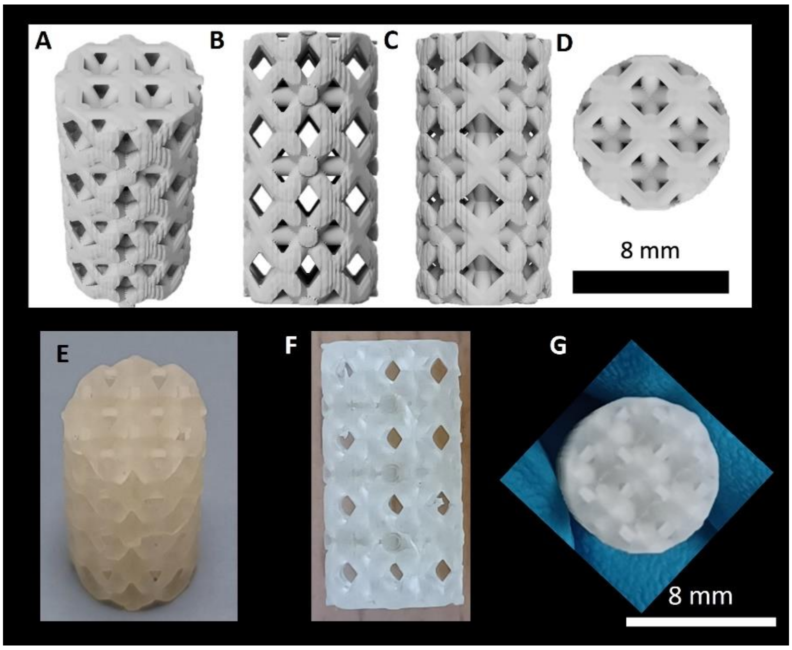


**S. Figure 1:** CAD model of the scaffold showing different views of the scaffold geometry. (A) Scaffold with intricate network of struts. (B) Diamond shaped pores within the scaffold. (C) The cross shape pattern gives strength with smaller pores between the internal struts. (D) The end of the scaffold with a cross-based configuration. (E) The scaffold printed was identical to that designed with the software. (F) The pores of the scaffold are clearly visible. (G) The end of the scaffold printed per the design. Scale bar 8 mm.

#### Method for coating PCL-TMA900 scaffolds with Laponite and BMP-2

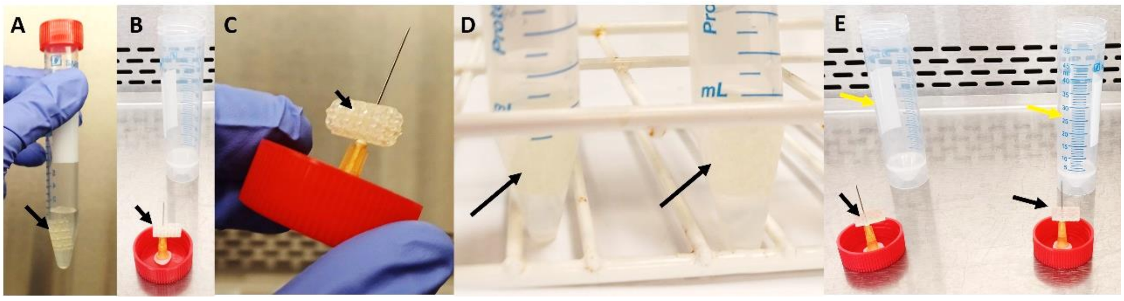


**S. Figure 2:** (A) The scaffold (black arrow) immersed in 1% Laponite. (B) The scaffold (black arrow) drying on a 25-gauge needle in the sterile class II MSC. (C) The Laponite (black arrow) could be seen as shiny clear areas between the pores as a dry film after 3 hours minimum drying time. (D) The scaffolds (black arrows) within low protein-binding tubes with BMP-2/PBS solution. (E) The scaffolds (black arrows) drying in the class II MSC and the bases of the tubes (orange arrows) were retained to contain the scaffolds for transport and to maintain sterility of the scaffolds.

### Investigation of the volume and location of Laponite coating of PCL-TMA900 scaffolds using radiodense barium sulphate

To investigate where and what volume of Laponite may adhere to the PCL 900 scaffolds for the sheep study, two studies were performed. The Laponite (1%) or $dH_20$ had Barium sulphate 98% w/w powder (E-Z-HD, EZEM, UK) added by addition of 0.52 g barium sulphate in 1 mL of $dH_20$ to make a suspension. 300 µL of this barium suspension was added to 6 mL of Laponite or $dH_20$. 2 mL of each solution was used to coat the scaffolds for 1 hour at RT. The scaffolds were then allowed to dry for 3 hours. The scaffolds were scanned using the Milabs µCT scanner and again after rinsing in 10 mL $dH_20$ in 10 mL falcon tubes on the tube rotator at 30 rpm. Studies confirmed the ability for Laponite to stay adhered to the scaffold visualised when labelled with barium sulphate solution compared to a scaffold which had no coating, as $dH_20$ would not adhere to the scaffold surface to hold the barium to the scaffold surface (**Supplementary Figure 3**).

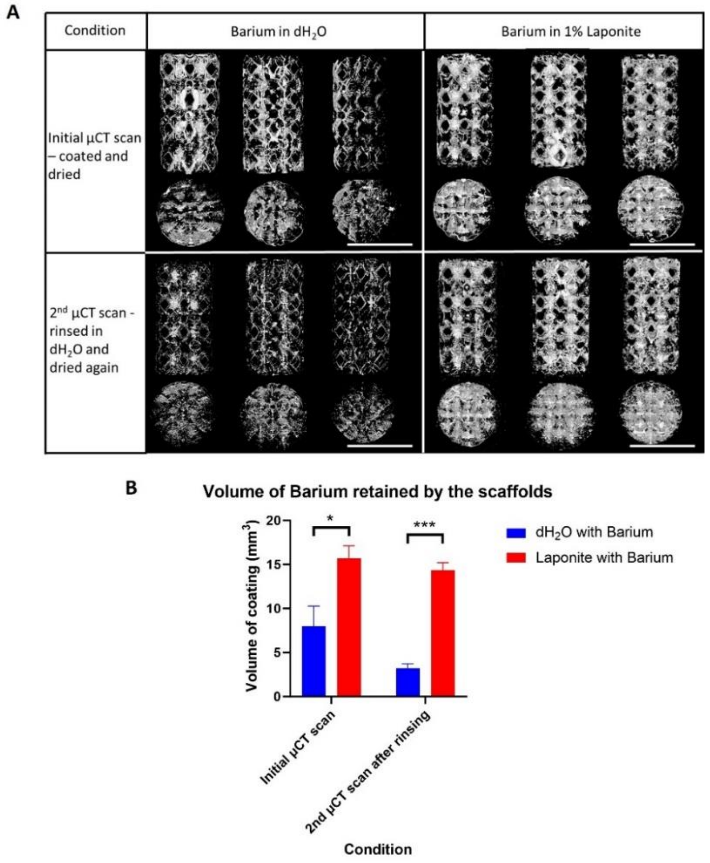

**S. Figure 3:** Determination of the location and persistence of Laponite adhered to the scaffold. (A) µCT images in the top row of PCL 900 scaffolds coated in $dH_2O$ with Barium or 1% Laponite with Barium showing the solution dry all over the PCL 900 scaffolds, especially in the Laponite coated group. Once rinsed, dried and re-imaged the scaffold retained more Laponite with Barium than $dH_2O$ with barium. Scale bar 8 mm. (B) Quantification of the radiodense barium on each scaffold showing greater persistence of Barium when held to the scaffold in Laponite than in $dH_20$. N=3 for each condition as shown. Mean and S.D. shown. Data analysed by multiple unpaired t-tests with Welch correction, *$p<0.05$, ***$p<0.001$.

### Optimising BMP-2 uptake by the PCL 900 scaffold

*Method of coating scaffolds with Laponite and BMP-2 for BMP-2 uptake ELISAs and a BMP-2 release study.*

The PCL-TMA900 scaffolds were coated as per the method described in the methods section and left to dry in a class II MSC before immersion in InductOS® BMP-2 (100 µg/mL or 200 µg/mL) diluted in PBS. The scaffolds were immersed in BMP-2 solution individually in low protein binding falcon tubes (Eppendorf) for 24 hours prior to removal and drying in a class II MSC. The residual BMP-2 solution was tested for the remaining BMP-2 concentration by ELISA.

*BMP-2 ELISA for quantification of BMP-2 remaining after PCL-TMA900 scaffold coating for in vitro and in vivo experiments.*

The PCL-TMA900 scaffolds were used for ELISA determination of BMP-2 binding at 100 µg/mL and 200µg/mL BMP-2 to the Laponite coated scaffolds prior to the study commencing, and with 100µg/mL BMP-2 for the sheep femoral condyle defect study.

The Quantikine® ELISA BMP-2 immunoassay kit was used to test the BMP-2 remaining in the scaffold coating solution, to deduce the quantity bound to the scaffold. Reagents and standard solutions were prepared as per kit instructions. The BMP-2 solutions were diluted in calibration buffer to enable results to be extrapolated from the standard curve. The large 8 × 15mm PCL 900 scaffolds were serially diluted from 100 µg/mL or 200 µg/mL to 1000 pg/mL using a minimum pipette volume of 10µL to reduce pipetting error. The standards were made as per kit instructions. 100 µL of assay diluent followed by 50 µL of standard, control, or sample was added to each well. The plate was shaken on a horizontal orbital microplate shaker at 450 rpm for 2 hours at RT. The solution was aspirated, and the wells washed and aspirated again 4 times. BMP-2 conjugate (200 µL) was added to each well and the incubation step on the shaker repeated for 2 hours. The aspiration and wash steps were repeated and 200 µL of substrate solution added to each well and the plate was left for 30 minutes at RT in the dark on the benchtop. Finally, 50 µL of stop solution was added to each well and the colour changed from blue to yellow. The plate was read using a GloMax® Discover microplate reader set to 450 nm. Results were calculated from the standard curve to determine the concentration of BMP-2 in each sample. In the sheep femoral condyle defect study, the 100 µg/mL stock coating solutions (with no scaffold immersed) were n=1, which was plated out in triplicate, while the 100 µg/mL solutions used to coat the Laponite coated scaffold samples were n = 8 and each plated out in triplicate and the average result of each scaffold triplicate was used for statistical analysis.

*Assessment of BMP-2 uptake from Laponite coated 8 × 15 mm PCL-TMA900 scaffolds.*

This experiment allowed assessment of BMP-2 uptake from larger scaffolds of approximately 10 times greater surface area than the octet-truss unit scaffolds used in previous studies (14) and at a higher concentration of BMP-2 in preparation for a large animal model study. The scaffold adsorbed approximately 40% of the BMP-2 from the initial 100 µg/mL coating solution, equating to at least 60 µg of BMP-2 taken up by the scaffold during coating.

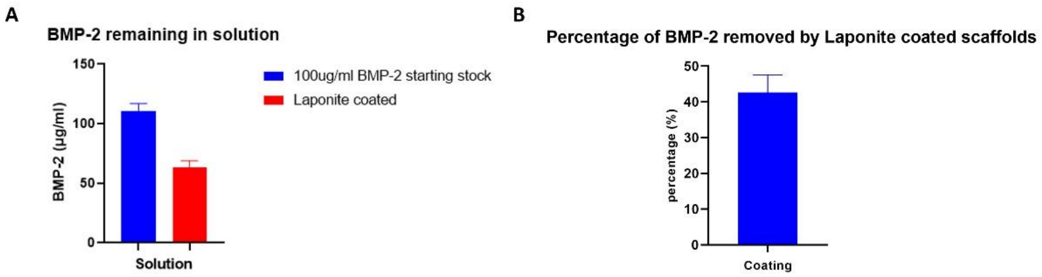


**S. Figure 4:** ELISA results of BMP-2 uptake from 3 large PCL-TMA900 scaffolds coated in Laponite. (A) The average concentration of BMP-2 remaining in the 3 eppendorf tubes used to coat the 3 large scaffolds. n=1 for stock solution with triplicate readings taken and n=3 for Laponite coated scaffolds with triplicate readings taken and averaged for each of the 3 samples, with the resulting 3 mean readings used to calculate mean and S.D. as shown. (B) The average percentage of BMP-2 taken up from the starting stock solution of 100 µg/mL was approximately 40%, n=3 with 3 readings taken form each scaffold and the 3 readings averaged. Mean and S.D. shown.

While ELISA results using 100 µg/mL coating solution confirmed BMP-2 binding to the scaffold of around 60 µg maximum from a total of approximately 150 µg of BMP-2 (i.e., approximately 40% binding), using 200 µg/mL gave approximately 20% binding (**S. Figure 5**), and therefore the scaffold was likely saturated with BMP-2 of a mass around 60 µg. Therefore, the limiting factor on BMP-2 uptake was ensuring adequate coating of the PCL-TMA900 scaffolds with Laponite.

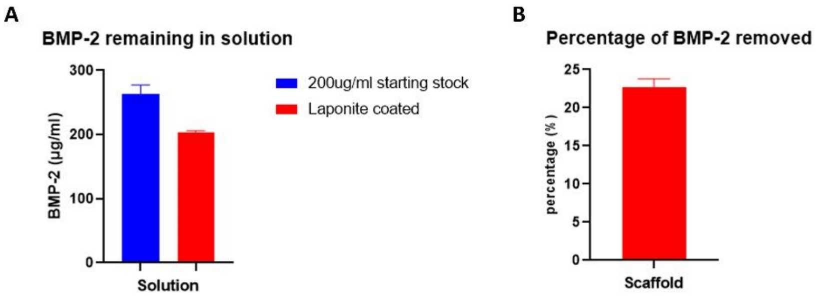


**S. Figure 5:** BMP-2 binding to Laponite coated scaffolds at 200 µg/mL concentration. (A) The quantity of BMP-2 remaining in solution after scaffold coating. N=3 for stock with triplicate readings from one sample and, n=2 scaffolds with readings taken from each in triplicate and averaged, and then the 2 averages displayed on the graph. Mean and S.D. shown. (B) The percentage of BMP-2 removed from the coating solution. N=2 scaffolds, with 3 readings from each of the 2 scaffolds and 3 readings averaged. Mean and S.D. shown.

*Method to assess BMP-2 release from Laponite/BMP-2 coated PCL-TMA900 scaffolds*

To try to elucidate the release mechanism of BMP-2 from the Laponite/BMP-2 coated scaffolds, experiments were performed in which C2C12 cells were used as a marker for BMP-2 presence in the media in which the scaffolds were incubated in DMEM. To determine if a bioactive quantity of BMP-2 was released from the PCL-TMA900 coated scaffolds when 100 µg/mL of BMP-2 was used, three scaffolds were coated with 2 mL 1% Laponite and allowed to dry, followed by 1.5 mL of 100 µg/mL BMP-2 and allowed to dry for 7 hours prior to being placed in 3 mL of DMEM/1% P/S in a 12 well plate. The media was collected into 15 mL low protein binding eppendorf tubes from each scaffold and replaced entirely. The timepoints assessed were 30 minutes, 2 hours, 4 hours, 6 hours, 24 hours. C2C12 cells were seeded into 24 well plates at $5\times10^4$ cells per well and cultured overnight in DMEM/1%P/S/10% FCS. The following day the media from each eppendorf for each time point was added to a well of C2C12 cells in biological triplicates and the cells cultured at 37°C for 2 days followed by ALP staining of the well.

*ALP staining of C2C12 cells cultured on tissue culture plastic*

Media was removed and the cells were washed twice in PBS followed by addition of 90% ethanol for 10 minutes, prior to washing again in PBS. Alkaline phosphatase activity was visualised by creating a solution of 4% (v/v) Naphthol AS-MX phosphate (Sigma) and 0.0024% (w/v) Fast Violet-B salt (Sigma) mixed in and adding 300µL staining solution to cells on TCP. Cells were incubated in the dark at 37°C for 1 hour. Red stain indicated positive ALP activity. $dH_2O$ was added to stop the reaction and the solution removed prior to imaging using a digital camera (Canon G10) attached to a stemi 2000-c stereomicroscope (Zeiss, UK).

*Assessment of BMP-2 release from Laponite coated PCL 900 8×15mm scaffolds*

Assessment of BMP-2 release from the 100 µg/mL BMP-2 coated PCL-TMA900 scaffolds was performed and the result confirmed BMP-2 release as C2C12 cells responded with marked red ALP staining until the 4 hour time point, showing less vibrant staining at the 6 hour time point (**S. Figure 6**). This continued response involved 3 changes of the media which may have washed off the surface BMP-2 which was not firmly held to the scaffold, however, if the BMP-2 was completely free of interaction with the Laponite coating the BMP-2 may have washed off at an earlier timepoint and staining may not have been as intense until the 4 hour timepoint.

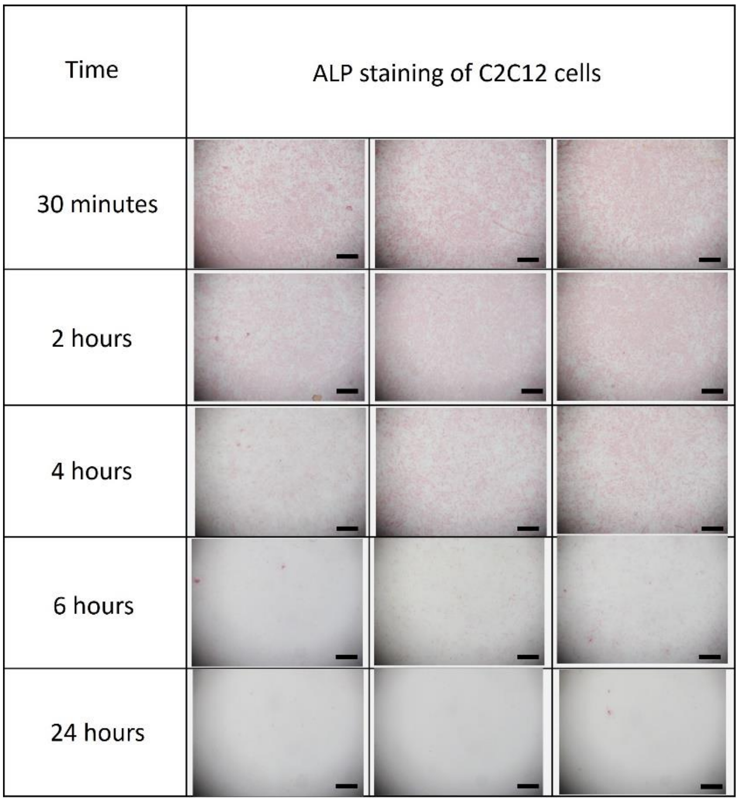


**S. Figure 6:** BMP-2 release assay using C2C12 cells to detect free BMP-2 in media. The C2C12 cells generated a positive red staining response until 6 hours, with most marked staining observed at 30 minutes, 2 and 4 hours. The BMP-2 was not found to be released at a detectable concentration over the following 18 hours, as the 24 hour time point showed no obvious red staining. N=3, scale bars 1 mm.

Therefore, in the sheep femoral condyle defect study 1% Laponite with 100 µg/mL of BMP-2 coating solution was used as there was no advantage to using BMP-2 solution of a higher concentration.

After the sheep study was complete, an ELISA revealed that an average ~50 µg of BMP-2 was on the PCL-TMA900 scaffolds implanted into the sheep. The sheep with the greatest BMP-2 uptake did not have the greatest mass of bone formed and conversely, the sheep with the least BMP-2 uptake did not have the least bone formed when the results were compared for each sheep. However, this was difficult to verify as if any Laponite was inadvertently removed from the scaffold then the BMP-2 would be bound to this rather than on the scaffold itself and the high concentration of BMP-2 in the PBS of pH 7 could lead to aggregation of the BMP-2 particles leading to some variability in ELISA results.

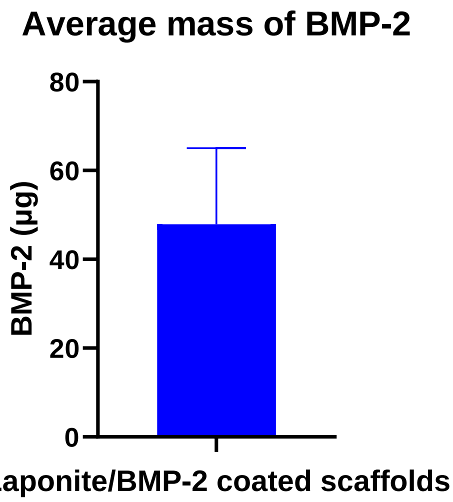


**S. Figure 7:** ELISA results of BMP-2 uptake on the scaffolds used in the sheep study. The mass of BMP-2 uptake varied in the sheep study with a mean mass of BMP-2 and standard deviation of 47.89 µg ± 17.13 µg. This was likely due to variation in Laponite mass and uneven coverage upon the scaffolds leading to variable BMP-2 binding. N=8, mean and S.D. shown.

### Sheep femoral condyle defect study layout

The sheep femoral condyle defect study was planned with n = 8 of each implantation material type allocated at random (Plain PCL 900, Laponite/BMP-2 coated PCL 900 scaffold, autograft or empty defect), therefore there were 32 defects. Due to the model being bilateral limb defects, 16 sheep were purchased for the study and group housed prior to use. The study plan is illustrated in **Supplementary Figure 8**.

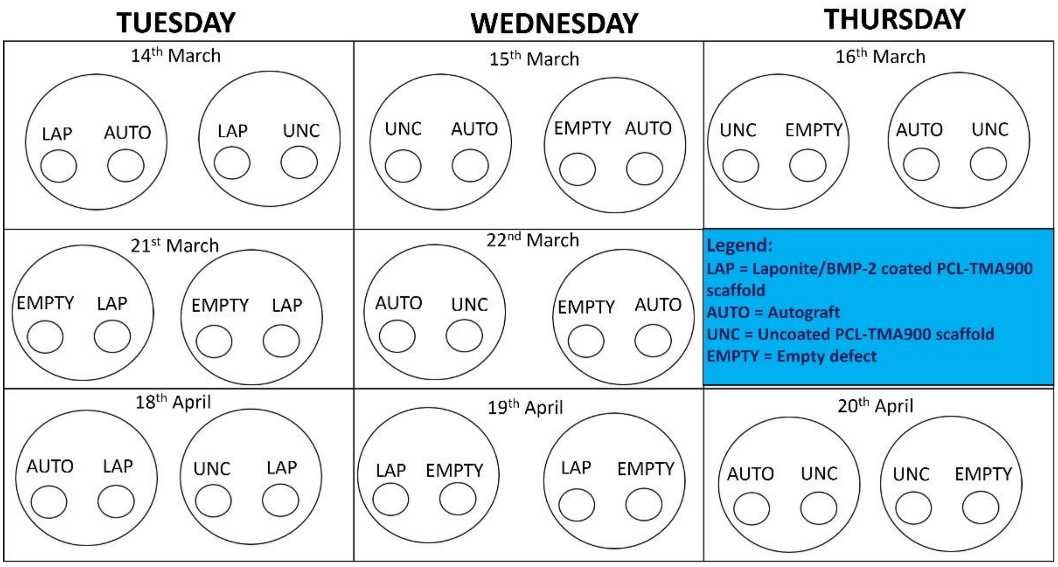

**S. Figure 8:** The sheep femoral condyle defect study plan showing the dates and materials to be implanted in each sheep. Dates included to enable comparison of seasonal bone formation in other studies to be compared. Each large circle represents a sheep with the left limb as the left smaller circle/label and right limb was the right smaller circle/label.

### Preparation for the sheep study – drill defect size and scaffold robustness

In preparation for the sheep study to assess the size of the scaffold for fit testing, an 8 mm diameter defect was drilled in the medial femoral condyle using the actual drill bit and drill to be used in the study and the 8 mm diameter x 15 mm tall scaffold inserted. The scaffold fitted exactly into the defect with no protrusion of the scaffold from the bone surface. In the actual study, the core of bone would be kept for use in autograft control defects in selected sheep (**Supplementary Figure 9**).

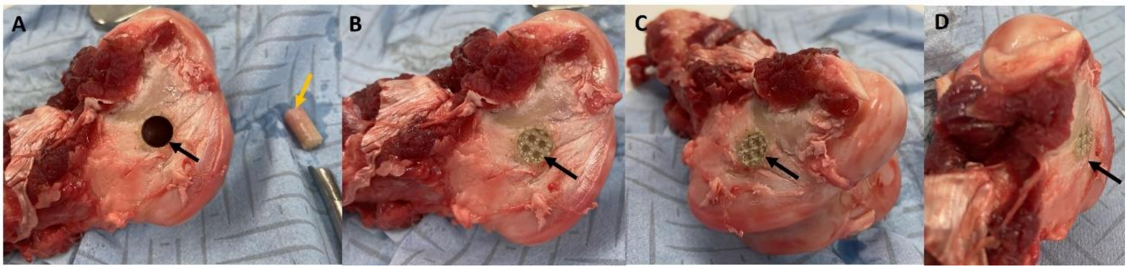


**S. Figure 9:** The medial femoral condyle bone defect with PCL 900 scaffold insertion. (A) The 8 mm × 15 mm drill bit was used to create a metaphyseal bone defect (black arrow) in the medial femoral condyle of the bone with the bone core (orange arrow) preserved for use in cases with autograft control implants. (B) The PCL 900 scaffold (black arrow) within the bone defect. (C and D) The PCL scaffold (black arrows) was in line with the bone surface with no protrusion or damage to the scaffold during implantation.

The Surgical procedure:

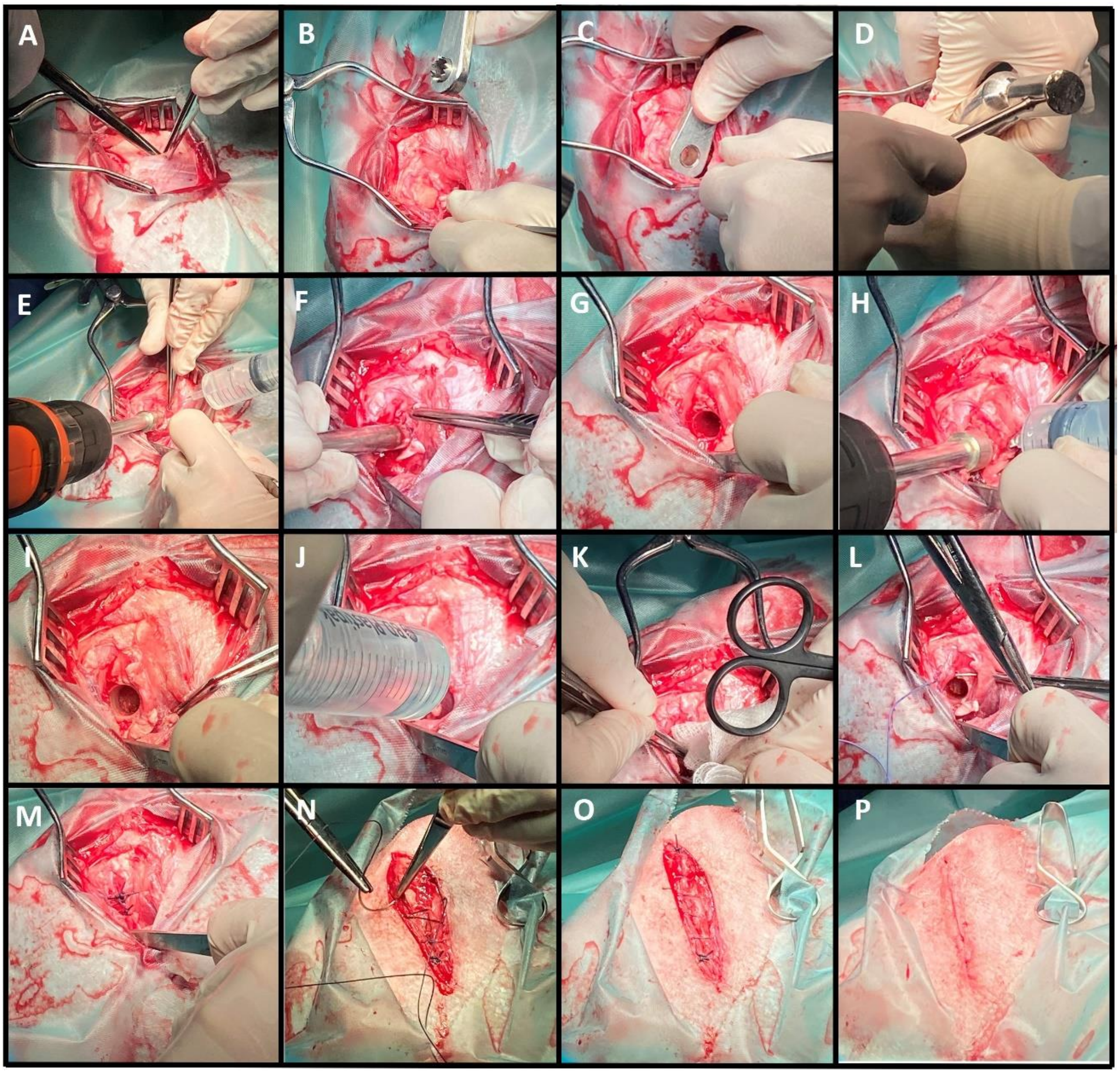


**S. Figure 10:** The surgical procedure: (A) The skin and underlying soft tissue over the medial femoral condyle was incised (B) Weitlaner retractors and a Hohmann retractor were used to aid visualisation of the medial femoral condyle. (C) A custom made guide was used to aid defect creation. (D) A mallet was used to create a seat for the drill bit using the guide (E) A single cylindrical drill hole/defect (8 mm diameter × 15 mm deep) was created in the cancellous bone region of the medial femoral condyle. (F and G) the bone core was removed using a specialised tool. (H and I) The defect was cleared of debris using a custom calibrated reamer to create a standardised defect. Throughout coring and reaming, the drills were cooled with sterile saline solution to prevent tissue damage. (J, K, L) The defect was lavaged with sterile saline to remove any bone fragments and dried using a sterile swab to check for bleeding, however none/very minor ooze was noted. (M) After implantation/defect creation, the periosteum was sutured over each defect/scaffold with absorbable suture (Vicryl 1-0, Ethicon, Kirkton, UK) which aided retention of autograft in the bone defect. (N and O) The subcutaneous tissue was closed using absorbable material (Vicryl 2-0, Ethicon, Kirkton, UK) in simple continuous suture pattern. (P) The skin was closed with absorbable material (Vicryl 2-0, Ethicon, Kirkton, UK) in an intradermal simple continuous suture pattern.

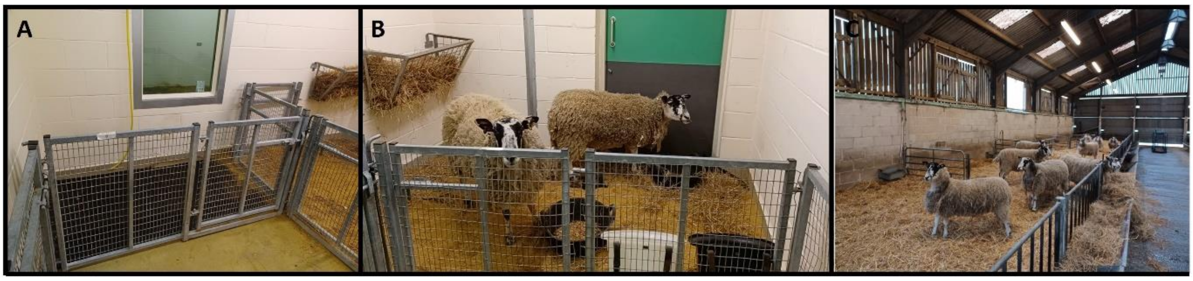


**S. Figure 11:** Recovery area for sheep and barn. (A) immediate recovery area; (B) pen for overnight holding of sheep who had surgery and, (C) the barn in which the sheep were housed during the study.

### µCT scan settings used for *in vivo* and *ex vivo* imaging

The scan information for acquisition and reconstruction parameters for the *in vivo* and *ex vivo* µCT imaging is shown in **Supplementary Table 1**.

**S. Table 1:** Settings used for µCT scanning of femoral condyles and trimmed samples.

| **Imaging reference** | **Whole medial condyle** | **Trimmed bone samples** |
|---|---|---|
| Voltage (kVp) | 55 | 50 |
| Current (mA) | 0.17 | 0.21 |
| Exposure time (ms) | 75 | 75 |
| Filter (µm thickness) | Al 400+100 | Al 400+100 |
| Scan angle (°) | 360 | 360 |
| Step angle (°) | 0.25 | 0.25 |
| Reconstruction voxel size ($\mu m^3$) | 40 | 20 |
| Pixel number | 40 | 20 |
| Frame averaging | 1 | 1 |

### Processing of bone tissue for µCT scanning

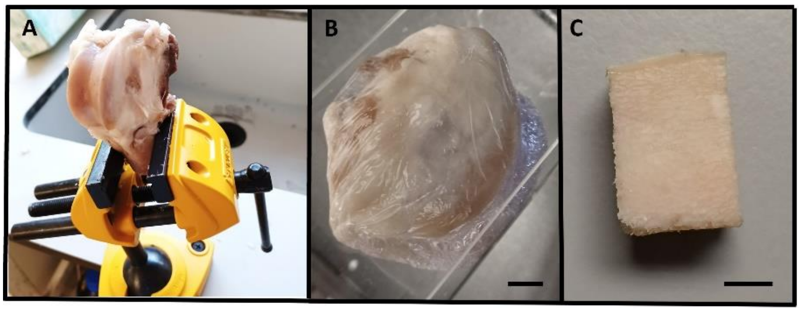


**S. Figure 12:** Processing the bone for µCT scanning: (A) trimmed condyle, (B) condyle in clingfilm for scanning scale bar 8 mm and, (C) Trimmed sample for scanning for higher resolution analysis, scale bar 8 mm.

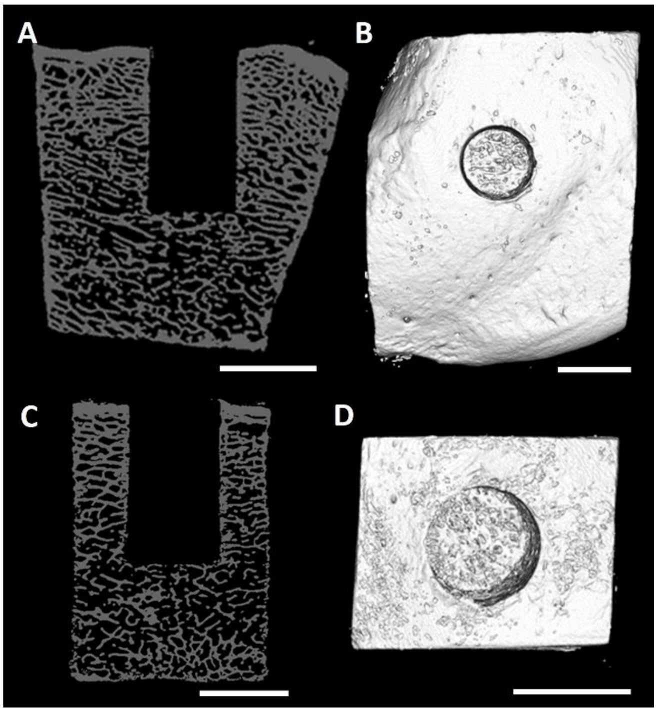


**S. Figure 13**: µct images of an empty defect. A cadaver sheep condyle had the defect hole created and µCT was used to illustrate the defect. (A) The empty defect in the medial condyle at 40 µm reconstruction voxel size, (B) A in 3D, (C) the empty defect in the trimmed condyle at 20 µm reconstruction voxel size, (D) C in 3D, scale bars 8 mm.

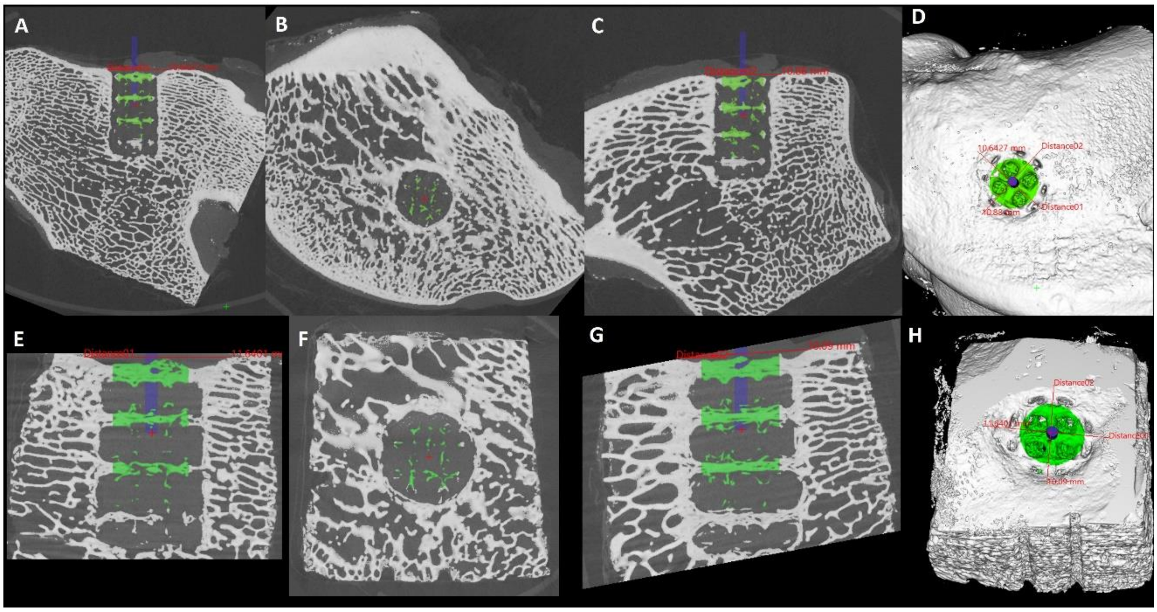


**S. Figure 14:** Example of the method of quantification of bone volume in the 6 mm x 12 mm defect volume within the centre of the 8 mm x 15 mm defect site. (A - D) 40 µm resolution images and (E - H) 20 µm resolution images. A line was drawn across the top of the defect from edge of cortex to edge of cortex in sagittal and coronal planes, and a blue line of 1 mm diameter × 12 mm length was drawn from the centre of this line/defect. This created a point at the end of the blue line exactly 6mm within the defect. The cursor was placed at the bottom of the blue line within the defect. A region of 6 mm diameter × 12 mm length was then created in green from this cursor to create a bone volume to be quantified from the top of the defect 12 mm deep × 6 mm wide.

### Processing of tissues for histology

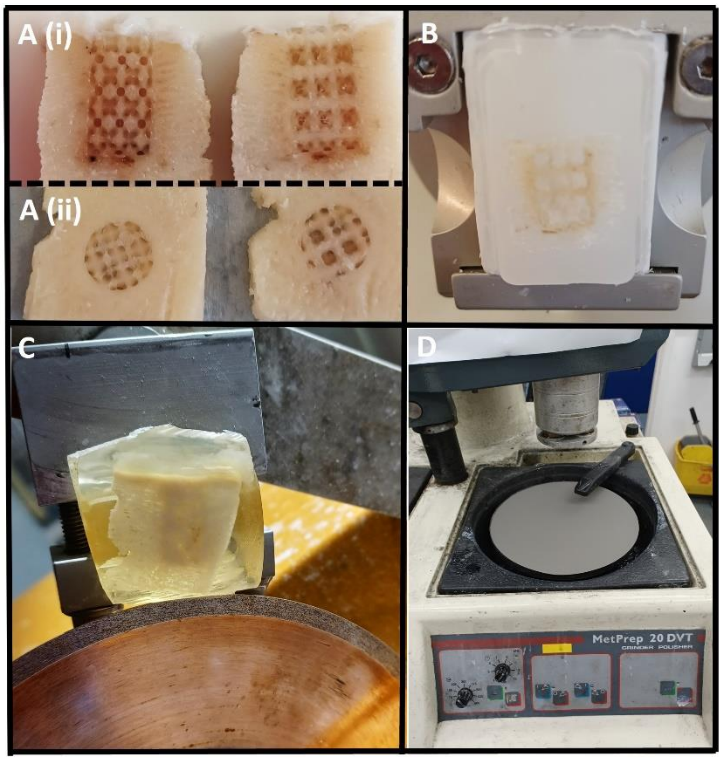


**S. Figure 15:** Processing the bone for histology. (A) Bone with a scaffold implant cut in half for resin and wax histology (i) longitudinally and (ii) transversely. (B) Bone and scaffold embedded in wax in the microtome for sectioning. (C) Bone and autograft within defect in resin being cut using the Isomet low speed saw on maximal speed of 10. (D) The MetPrep 20 DVT machine with the rotating plate with sandpaper and black water tap to remove debris and aid smooth grinding of resin embedded samples as thinly as possible prior to staining.

Further histology images demonstrating bone within the pores of the Laponite/BMP-2 coated scaffold compared to fibrous tissue in the empty and uncoated scaffold groups and irregular bone in the autograft group

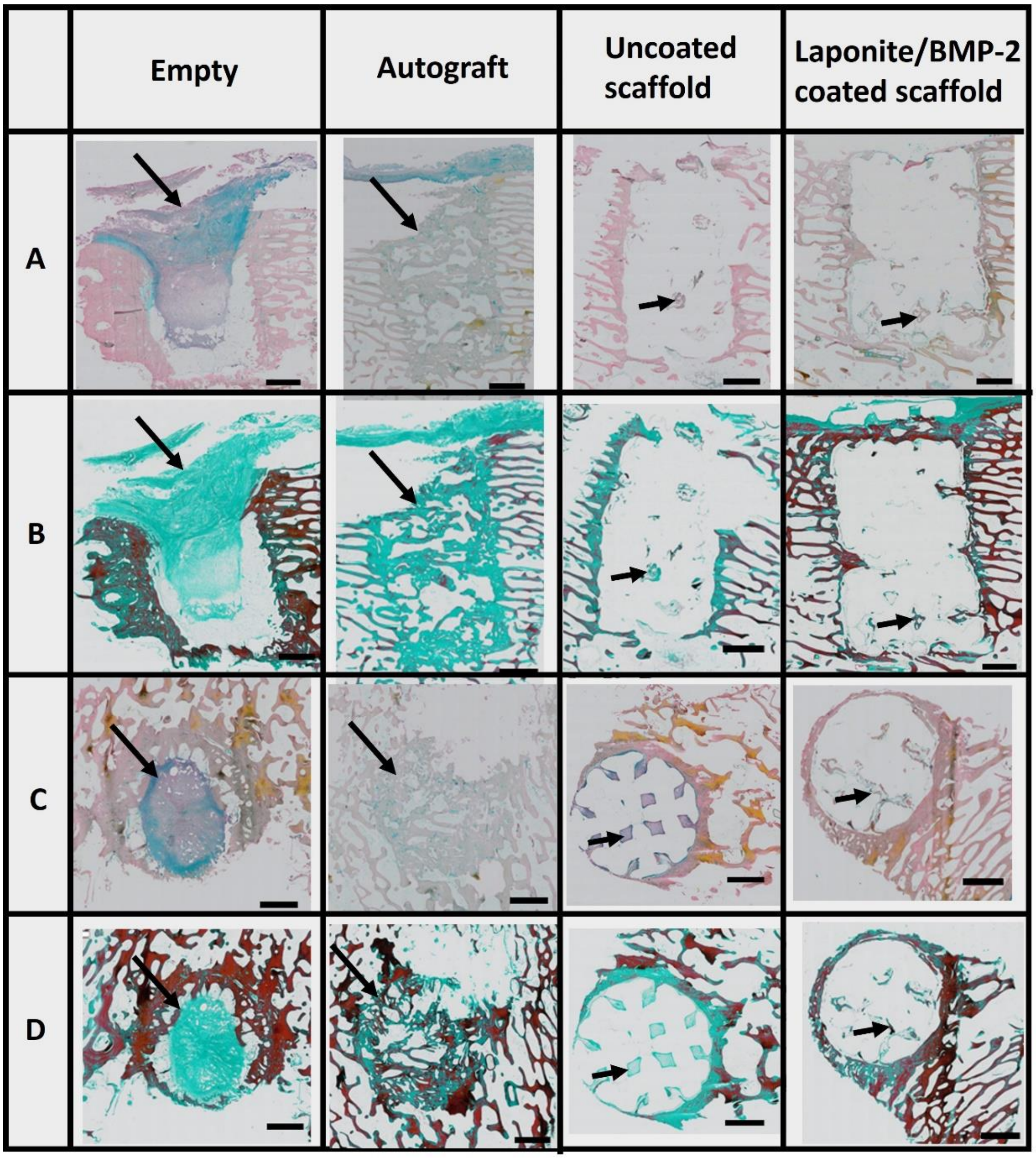


**Figure S. 16:** Representative images of the tissue formed within each defect group. Rows A and C are Alcian blue and Sirius red stained sections to illustrate proteoglycans (blue) and collagen fibres (red). Rows B and D are Goldner's trichrome stained sections to show the collagen tissue (green) and osteoid (red). The empty defect contained a plug of fibrous tissue (arrows) within the centre and top 2/3 of the defect with adipose tissue at the base of the defect. The autograft group contained bone (arrows) filling the defect with regions of marrow between the randomly positioned struts of bone in comparison to the parallel native bone seen around the periphery of the defect within the medial femoral condyle. The uncoated scaffold showed fibrous tissue within the pores of the scaffold (arrows), while the Laponite/BMP-2 coated scaffold formed bone and marrow tissue around the scaffold (arrows). Scale bar 2 mm.

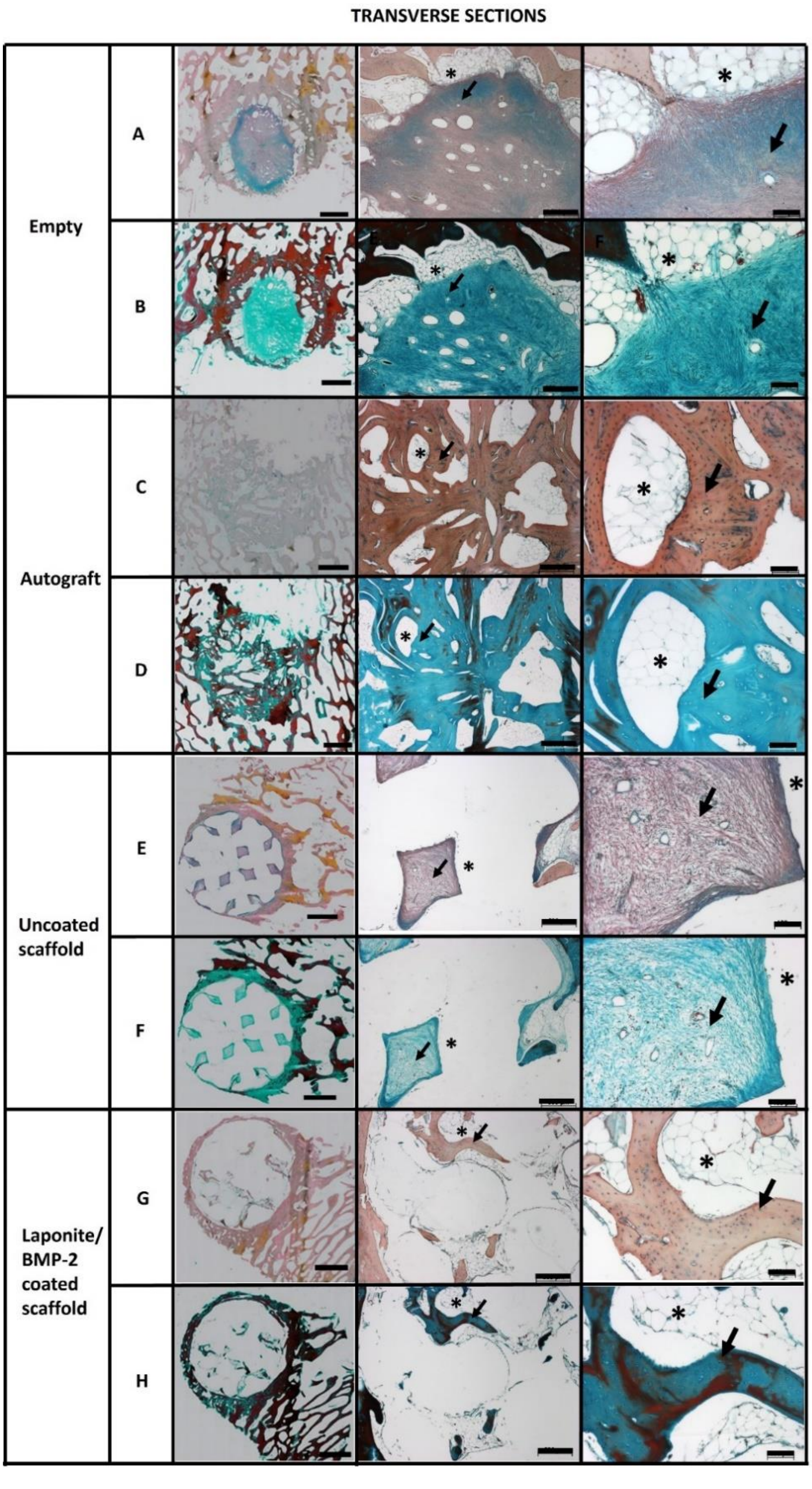


**Figure S. 17:** Representative images of histology of transverse sections of each defect type. (A, C, E, G) Alcian blue and Sirius red staining for proteoglycan and collagen respectively, (B, D, F, H) Goldners trichrome staining for collagen (green) and osteoid (red). First column are overview images of the defect in each group, scale bars 2 mm. Empty defect (rows A and B) with fibrous tissue (arrow) and marrow tissue (asterix). Autograft (rows C and D) with bone (arrow) and marrow (asterix). Uncoated scaffold (rows E and F) with fibrous tissue (arrow) in a diamond shape within the pores of the scaffold material (asterix) and Laponite/BMP-2 coated scaffold (rows G and H) with bone (arrow) and marrow tissue (asterix). 2nd column scale bars 500 µm and 3rd column scale bars 100 µm.

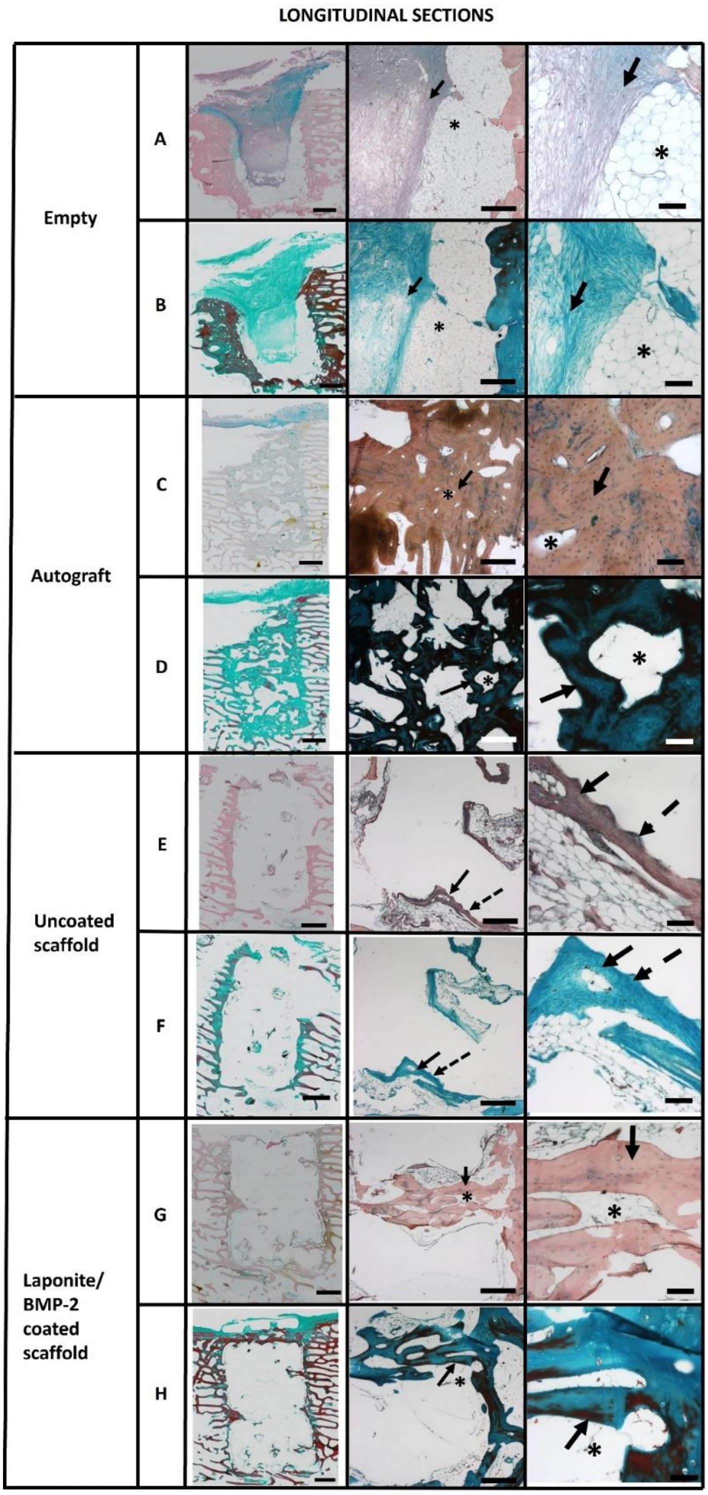


**Figure S. 18:** Representative image of histology of longitudinal sections of each defect type. (A, C, E, G) Alcian blue and Sirius red staining for proteoglycan and collagen respectively, (B, D, F, H) Goldners trichrome staining for collagen (green) and osteoid (red). First column are overview images of the defect in each group, scale bars 2 mm. Empty defect (rows A and B) with fibrous tissue (arrow) and interdigitation of tissue with the scaffold ridges (dashed arrow). Autograft (rows C and D) with bone (arrow) and marrow (asterix). Uncoated scaffold (rows E and F) with fibrous tissue (arrow) in a diamond shape within the pores of the scaffold material (asterix) and Laponite/BMP-2 coated scaffold (rows G and H) with bone (arrow) and marrow tissue (asterix). 2nd column scale bars 500 µm and 3rd column scale bars 100 µm.
